# Multivariate Neural Markers of Individual Differences in Thought Control Difficulties

**DOI:** 10.1101/2025.02.04.636283

**Authors:** Jacob DeRosa, Harry R. Smolker, Hyojeong Kim, Boman Groff, Jarrod Lewis- Peacock, Marie T. Banich

**Affiliations:** Department of Psychology and Neuroscience, University of Colorado Boulder; Institute of Cognitive Science, University of Colorado Boulder; Department of Psychology, University of Texas at Austin

**Keywords:** Individual Differences, Working Memory, Cognitive Control, Multi-voxel Pattern Analysis, Representational Similarity Analysis, Cortical Gradients, Functional Connectivity, Default Mode Network, Frontoparietal Network

## Abstract

Difficulties in controlling thought, including pathological rumination, worry, and intrusive thoughts, occur in a range of mental health disorders. Here we identify specific patterns of brain activity distributed within and across canonical brain networks that are associated with self-reported difficulties in controlling one’s thoughts. These activity patterns were derived using multivariate pattern analysis on fMRI data recorded while participants engaged in one of four operations on an item in working memory: maintaining it, replacing it with another, specifically suppressing it, or clearing the mind of all thought. Individuals who reported greater difficulties exhibited brain activation patterns that were more variable and less differentiated across the four operations in frontoparietal and default mode networks, and showed less distinct patterns of connectivity within the default mode network. These activity profiles were absent during rest but serve as promising task-based neural markers, explaining over 30% of the variance in thought control difficulties.

## Main

The goal of the present paper is to examine what aspects of neural function may be associated with self-reported individual differences in the ability to control thoughts. This question is of importance as many different types of psychopathology are characterized by repetitive negative thinking (e.g., rumination and worry) that is often difficult to control and associated with psychological distress. For instance, the level of such repetitive negative thinking is predictive of the subsequent severity and length of depressive episodes (Susan Nolen-Hoeksema et al., 2008). While the associations between such thought patterns and psychopathology is clear, less is known about the neural mechanisms that might underlie such repetitive negative thoughts. In particular, it is of interest to examine what cognitive and neural mechanisms might allow such thoughts to be expunged from the current focus of attention in working memory (WM), which is a buffer that enables us to transiently hold online 4-7 items for a relatively short time (e.g., up to ∼10 seconds) (Cowan, 2011; Oberauer, 2019). While much research has examined how information is gated into WM (Badre, 2012; Kessler, 2017) or how attention is shifted to different pieces of information in WM (Ku, 2018; Oberauer, 2019), up until now, it has been challenging to both verify that an item has been removed from WM and to characterize the neural mechanisms that support such removal operations. Recently, Kim et al. (2020) used neuroimaging and multi-voxel pattern analysis (MVPA) to track the removal of information from WM and to build on prior work that characterizes the operations that allow for such removal (Banich et al., 2015). In particular, this work demonstrated that information can be removed from WM in at least three distinct ways: by replacing an item with something else, by specifically suppressing it, and by clearing the mind of all thought. Moreover, univariate analyses, as well as multivariate classifier analyses, have shown that these operations are distinct, each with its own neural pattern (Banich et al., 2015; Kim et al., 2020).

In the present study, we investigate the degree to which properties of these neural patterns may be related to difficulties in removing information from WM. This research question is motivated, in part, by prior studies showing that individuals with high levels of rumination and/or worry exhibit alterations in patterns of neural activation and connectivity (Kertz et al., 2015; Satyshur et al., 2018; Sokołowski et al., 2022). We focus on processes relevant to rumination and worry because they manifest as difficulties in controlling the contents of WM, as compared to intrusive thoughts that characterize post-traumatic stress disorder (PTSD), which are associated with difficulties in suppressing the retrieval of information from long-term memory (Catarino et al., 2015; Küpper et al., 2014), which has been shown to be associated with cognitive control over hippocampal regions (Anderson & Hulbert, 2021).

We also examine the hypothesis that brain activation patterns during the WM operations discussed above (maintain, replace, suppress, clear) are associated with self-reported difficulties in controlling thoughts in three distinct manners. First, we examine whether such difficulties are associated with univariate activation in regions associated with these control operations. Second, we examine whether multi-voxel patterns of activation vary as a function of an individual’s difficulty in controlling their thoughts, which, to preview, yielded multiple intriguing associations. Third, we explored whether connectivity between brain regions that implement these control operations varies as a function of an individual’s difficulty in controlling their thoughts.

With regards to associations with multi-voxel patterns of activation, we examine various manners in which self-reported difficulties in controlling thoughts might manifest neurally. One possibility we explore is that individuals with difficulty controlling thoughts have poorly differentiated neural mechanisms for implementing control operations over information in WM. We investigate this possibility in two manners, one using an MVPA classifier approach and another using representational similarity analysis (RSA).

Our prior work has shown that MVPA classifiers, which use machine learning algorithms, can effectively categorize a neural pattern on a given trial as belonging to one of the four WM operations (maintain, replace, clear, suppress), and that there is a fair amount of consistency of these neural signatures across individuals (Bretton et al., 2024; Kim et al., 2020). Said differently, the MVPA classifier can correctly predict the operation which an individual was instructed to perform based on the pattern of neural activation. Here, we examine the hypothesis that individuals with difficulty controlling thoughts have poorer or less “crisp” representations of these operations, meaning they are less differentiable at the neural level, which would be reflected in poorer accuracy of the MVPA classifier to predict the actual operation in which an individual was engaged.

In the second approach, we ask the same question of whether the representations of these operations are less differentiable from each other in individuals with difficulties in controlling their thoughts, but using RSA, which examines patterns of activation across brain regions, but without trying to classify them into a specific category. Recent work re-analyzing data from Kim et al. (2020) used RSA in a novel manner to identify four distinct brain networks (DeRosa et al., 2024), which we refer to as _wmo_networks (_wmo:_ working memory operation). Each of these networks consists of a set of brain regions (specifically parcels drawn from Glasser et al., 2016) that all represent a given WM operation (i.e., maintain, replace, clear, suppress) in a similar manner, as reflected in similar activation patterns across brain regions for a given operation. However, the configuration of those multi-voxel patterns across the four operations differs across the _wmo_networks.

The four representational _wmo_networks identified by DeRosa et al. (2024) are referred to by the brain regions with which they most overlap: the Visual _wmo_Network (V_wmo_), Somatomotor _wmo_Network (SM_wmo_), Default Mode _wmo_Network (DM_wmo_), and Frontoparietal Control _wmo_Network (FPC_wmo_). These _wmo_networks are distinct from intrinsic connectivity networks (e.g., Yeo-7 network parcellation (Yeo et al., 2011)) that represent regions whose activation profiles co-vary over time. A figure illustrating these neural representation patterns across the V_wmo_, SM_wmo_, DM_wmo_, and FPC_wmo_ can be found in **Supplemental Figure 1**. Within the V_wmo_, the maintain and replace operations have similar neural representations (i.e., multi-voxel patterns), distinct from suppress and clear, which are also similar to each other. In the SM_wmo_, clear is represented distinctly from the other three operations, which are represented similarly. In the DM_wmo_, maintain and replace are represented similarly but distinct from suppress and clear, which are also distinguishable from each other. Finally, the FPC_wmo_ has distinct representations for each of the four operations. Here, we examine the hypothesis that the individuals with higher levels of difficulty in controlling thoughts have less differentiable representations of the working operations in that their average multi-voxel pattern across a given _wmo_network is less distinguishable from the median pattern across all networks than is observed for individuals who report lower levels of difficulty.

A second possibility, not mutually exclusive with the first, which is afforded by performing RSA in these _wmo_networks, is that the neural implementation of these removal operations *within* a given network (as compared to between them) is more variable across the constituent parts of that network. Said differently, we propose that difficulties in controlling thoughts may be associated with representations of the WM operations that are less consistent or concordant within a given _wmo_network.

Finally, we examined whether difficulties in controlling thoughts might also be associated with altered connectivity between the constituent parts of these _wmo_networks. This question is of interest because previous research has found that difficulties in worry, repetitive negative thinking, and rumination are associated with altered connectivity within and between neural networks (Ehring et al., 2011; Hamilton et al., 2011; Li et al., 2016). Hence, here we test the hypothesis that higher levels of difficulties in controlling thoughts will be associated with reduced connectivity between the subregions within each of the DeRosa et al. (2024) _wmo_networks discussed above (V_wmo_, SM_wmo_, DM_wmo_, and FPC_wmo_).

To evaluate the meaningfulness of any associations we might observe between difficulties in controlling thoughts and these _wmo_network metrics, we also tested the degree to which these alterations are specific to situations in which individuals are actively performing WM-control operations. Suppose we find, as hypothesized, that individuals with difficulty controlling their thoughts have neural representations that are less differentiated between networks or are more variable within networks or that they have reduced connectivity within networks. In each case, it is important to determine whether these organizational characteristics are specific to WM-control operations or whether they reflect a more general pattern of brain disorganization that might exist across most or all mental states (Greene et al., 2018). To do so, we examined whether brain activation patterns at rest can predict an individual’s level of difficulty in controlling thought. If the neural patterns observed during the WM operations can predict thought difficulties, but those at rest cannot, it would provide supportive evidence that these neural patterns are specific to the engagement of thought control processes and not a general property of an individual’s brain organization.

In sum, we investigate the hypothesis, using complementary MVPA approaches, that self-reported difficulties in controlling information in WM are associated with alterations in neural activity during cognitive operations to exert control over the contents of WM.

## Methods

All neuroimaging analyses in the current study utilize neuroimaging data initially reported by Kim et al. (2020) and re-analyzed in a novel manner by De Rosa et al. (2024). Complete data collection procedures are described in Kim et al. (2020), with essential details just highlighted below. Some analyses in the present utilize data from those two publications, while other analyses are novel to the present report. In the methods below, we differentiate between these types of data. Furthermore, while collected as part of the study initially reported by Kim et al., 2020, new to the current report are questionnaire data used to assess differences among individuals in their ability to control their thoughts.

### Participants

The current study involves a total of 48 participants (16 male; age, M=23.65, SD=5.06), which is one less than the 49 reported in Kim et al. (2020). One participant was excluded from the Kim et al., 2020 study due to fMRI classifier performance below chance level (area under the Receiver Operating Characteristic (AUC) curve < 0.5). All participants had normal or corrected-to-normal vision, provided informed consent, and were compensated $75. The study was approved by the University of Colorado Boulder Institutional Review Board (IRB protocol # 16-0249).

### Behavioral Measures

To derive a measure that would capture an individual’s difficulty in controlling their thoughts, we created a composite measure called “thought control difficulties”. This measure was created by averaging the z-scores from three scales: 1) the White Bear Suppression Index (WBSI) (Wegner & Zanakos, 1994); 2) The Penn State Worry Questionnaire (PSWQ) (Meyer et al., 1990); and 3) the Brooding subscale score from the Ruminative Response Scale (S. Nolen-Hoeksema et al., 1999). We used an average of these Z-scored measures because we (Groff et al, unpublished data) have found in a larger sample of 1007 individuals that the average of these measures is highly correlated (r=.0.95) with a common factor underlying these three measures derived from a confirmatory factor analysis (CFA; see **Supplemental Materials**).

Refer to **Figure 1** for the correlations of these three subscales in the current sample and the thought control difficulties score.

**Figure 1.**
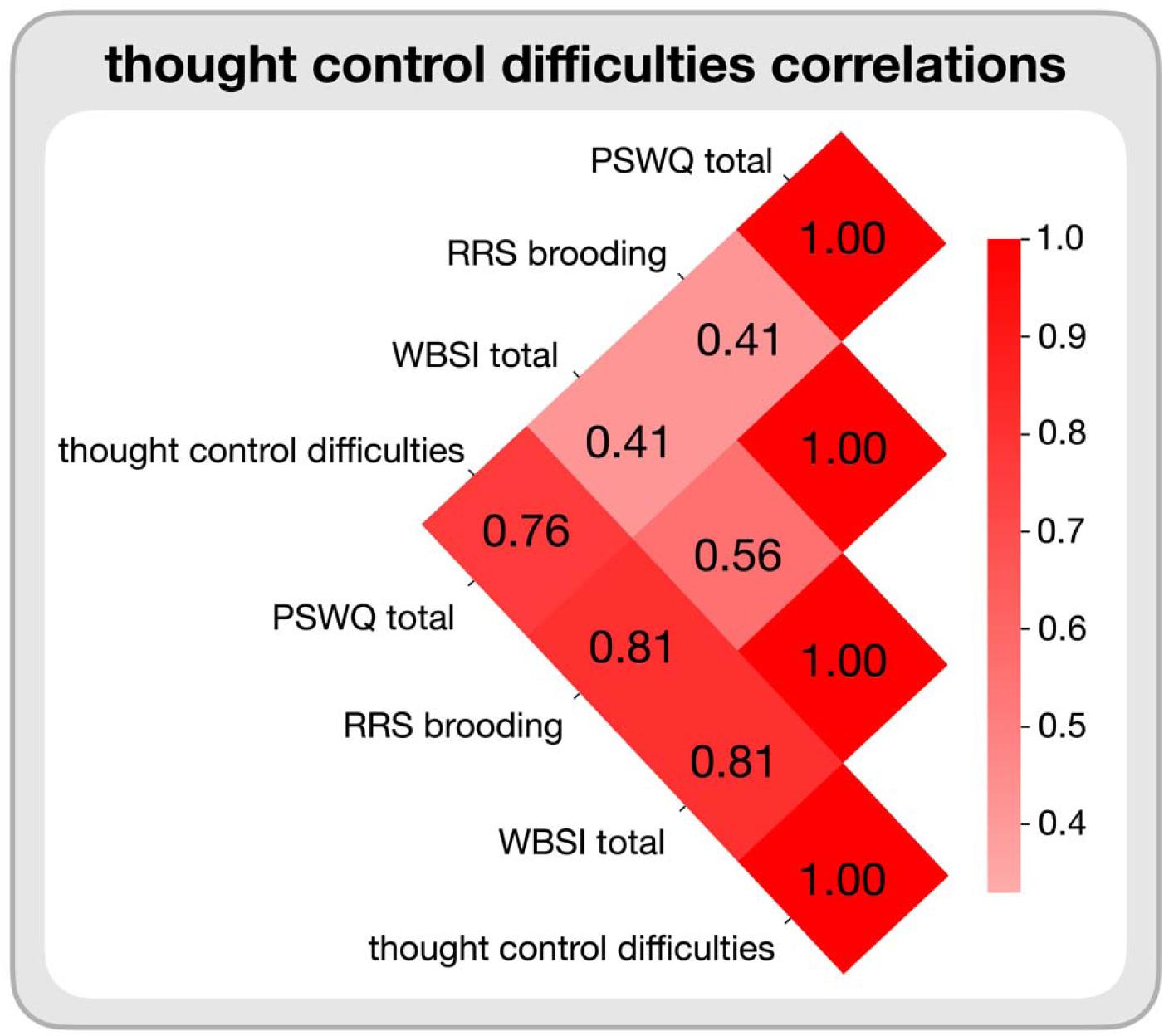
Pearson correlations between the PSWQ, WBSI, and RRS brooding subscale scores and the thought control difficulties score were calculated by taking the Z-score average of the PSWQ, WBSI, and RRS brooding subscale scores.

The WBSI is a 15-item self-report scale, with items rated from 1 (strongly disagree) to 5 (strongly agree). It includes two primary subscales: Suppression and Intrusion. Sample item from the Suppression subscale include “There are things I prefer not to think about” and “I have thoughts I try to avoid.” The Intrusion subscale features items such as “I have thoughts that I cannot stop” and “There are thoughts that keep jumping into my head.” The PSWQ measures the tendency to worry excessively and uncontrollably. It comprises 16 items scored on a 5-point Likert scale ranging from 1 (not at all typical of me) to 5 (very typical of me). This scale include both regularly scored items, such as “I know I should not worry about things, but I just can’t help it,” and reverse-scored items, like “I don’t tend to worry about things.” The brooding subscale from the RRS includes 5 items, measuring the extent to which an individual engages in repetitive focus on the causes and consequences of distress, with an example question being: “Why can’t I handle things better?”. It is scored on a scale of 1 (“almost never”) to 4 (“almost always”).

### Experimental Design

#### Stimuli

Stimuli for the study consisted of colored images (920 × 920 pixels) from three categories with three subcategories each: faces (actors, musicians, politician), fruit (apples, grapes, pears), and scenes (beaches, bridges, mountains). Faces were recognizable celebrities, and scenes were recognizable locales (e.g., a tropical beach) or famous landmarks (e,g, the Golden Gate Bridge). Images were obtained from various resources, including the Bank of Standardized Stimuli and Google Images. Six images from each subcategory were used, for 18 images per category and 54 images in total.

#### Data Acquisition

MRI data were acquired on a Siemens PRISMA 3.0 Tesla scanner at the Colorado University at Boulder Intermountain Neuroimaging Consortium Core Facility (RRID:SCR_018990). Structural scans were acquired with a T1-weighted sequence, with the following parameters: repetition time (TR) = 2400 ms, echo time (TE) = 2.07 ms, field of view (FOV) = 256 mm, with a 0.8 × 0.8 × 0.8 mm^3^ voxel size, acquired across 224 coronal slices. Functional MRI (fMRI) scans were obtained using a sequence with the following parameters: TR = 460 ms, TE = 27.2 ms, FOV = 248 mm, multiband acceleration factor = 8, with a 3 × 3 × 3 mm^3^ voxel size, acquired across 56 axial slices and aligned along the anterior commissure-posterior commissure line.

#### fMRI Procedure

The experimental aspects of the study consisted of two phases: a functional localizer and the central study. The data used in the present report were drawn from the resting-state data and from the central study which was designed to allow the track the representational status of an item in WM item while it was being manipulated using one of four distinct cognitive operations: maintaining an image in WM (maintain), replacing an image in WM with the memory of an image from a different category (e.g., replacing an actor with an apple), suppressing an image in WM (suppress), and clearing the mind of all thought (clear).

On each trial, participants were presented with an image for 6 TRs (2760 ms) followed by another 6 TRs of an operation screen instructing participants how to manipulate the item in WM, and then a jittered inter-trial fixation lasting between 5 and 9 TRs (2300–4140 ms), consisting of a white fixation cross centered over a black background. The operation screen consisted of two words in the top and bottom halves of the screen, presented over a black background. For the maintain, suppress, and clear operations, the two words were the same: maintain, suppress, or clear, respectively. In the replace conditions, the word in the top half was switch, whereas the word in the bottom half indicated the image that the participant should switch to (e.g., apple). Participants were instructed to only switch to thinking about an image that had previously been shown during the functional localizer task.

Each image exemplar appeared at least once per operation condition across the entirety of the task. Trials were ordered pseudo-randomly within runs, with the order of trials optimized for BOLD deconvolution using optseq2 (Dale, 1999). Hence there were a total of 288 trials, with 72 trials for each operation.

Resting-state data were collected at the start of the fMRI session, prior to the functional localizer and central study, to provide baseline measures of neural activity. Participants completed two resting-state runs, each lasting approximately 5 minutes and 45 seconds. With a TR of 0.46 seconds, each run consisted of 750 TRs, resulting in a total of 1500 TRs across the two runs. During resting-state scans, participants were instructed to remain still, keep their eyes open, and fixate on a white cross displayed on a black background.

### Statistical Analyses

#### Univariate fMRI analyses

The univariate analyses were minimally examined and reported by Kim et al. (2020) (**Supplementary Figure 1**) and are expanded upon in the present report. Standard fMRI preprocessing processing procedures were carried out using the FSL suite (version 5.0.10) (http://fsl.fmrib.ox.ac.uk) including motion correction via ICA-AROMA (version 0.3 beta), high-pass filtering (100 s), and BET brain extraction. Registration of EPI images into subject- and standard-spaces was executed using FLIRT (version 6.0). Individual subject EPI images were registered to that subject’s MPRAGE structural image via linear Boundary-Based Registration and then registered to the MNI-152 template via 12 degrees of freedom linear transformation. The resulting EPI images were smoothed using an 8 mm full-width half-maximum Gaussian smoothing kernel. FEAT (version 6.00) was used to model effects during the 6 TRs (2760 ms) during which the manipulation of the item was occurring, with fixation blocks at the beginning and end of each run serving as a baseline. The TRs during the presentation time of the stimulus and the TRs during the inter-trial interval fixations both served as EVs of no interest.

Here, we focused on more contrasts than reported in Kim et al. (2020). They focused on broad contrasts such as manipulation vs. maintenance (replace, suppress, clear > maintain), removal vs. retention (suppress, clear > maintain, replace), and clearing all vs. other operations (clear > maintain, replace, suppress) as was done by Banich et al., (2015). In our study, we expanded on these by including 16 more specific contrasts between operations (e.g., maintain vs. replace, suppress vs. clear, and replace vs. clear, etc.) to investigate neural activation patterns across the WM-control operations. We examined how brain activation varied as a function of individual differences in self-reported difficulty in controlling thoughts. Thought control difficulty scores were included as a covariate in FEAT to evaluate their relationship with neural activation patterns. This allowed us to identify specific brain regions where activation levels during WM-control operations were associated (either positively or negatively) with individual differences in thought control abilities. Group-level significance testing was conducted using permutation-based non-parametric testing via FSL’s randomise tool, with threshold-free cluster enhancement (TFCE) applied to control for multiple comparisons and a family-wise error (FWE) corrected significance threshold of *p* < 0.05.

#### Accuracy of Whole-Brain Classifiers of WM-Control Operations

Here we utilized, for each participant, the already calculated accuracy with which the classifier distinguished between multi-voxel activation patterns for the four WM-control operations, as described in Kim et al. (2020). This accuracy score was based on the proportion of correctly classified operations across 288 trials (72 per operation). Kim et al. (2020) used a one-vs-other approach, where each WM-control operation was classified against the other three using multivariate pattern analysis (MVPA) with L2-regularized logistic regression. Higher accuracy scores indicate that the neural activation patterns corresponding to each operation are more distinct from the other 3 operations, reflecting an individual’s ability to reliably differentiate and implement the WM-control operation across trials.

### Representational Similarity Analysis

#### Analysis during working memory operations

The data for this analysis were drawn from De Rosa et al. (2024) which provides details of how we performed the RSA to identify the _wmo_networks. A summary figure of the approach is provided in **Supplemental Figure 1**. For this study, we used the individual RSA patterns from DeRosa et al. (2024), which were generated separately for each participant and each of the four WM-control operations. Specifically, for each individual and WM-control operation, the Glasser parcellation (Glasser et al., 2016) was converted from MNI space to native space, and the multivoxel activation patterns were extracted for each parcel in the native space. For each trial, the activation patterns were averaged across the operation period (i.e., 2.75 s, 6 TRs) from the onset that was shifted forward 4.6 s (10 TRs) to account for hemodynamic lag. The averaged patterns were weighted using the beta estimate contrast, a target-versus-non-targets t-contrast (e.g., face vs. {fruit, scene}) computed using SPM12 to identify category-selective voxels within the ventral visual stream ROI. In the GLM model, four operations utilizing a canonical hemodynamic response function and six motion regressors were modeled. Each operation weight was obtained from the t-contrast (e.g., *maintain* > others) without thresholding. The similarity of the weighted patterns across trials (i.e., 288 trial vectors total, 72 trials/operation) were then computed with Pearson’s correlation (i.e., RSA) to create the operation similarity matrix (288 x 288) for each parcel. The details of how we performed the RSA to identify the _wmo_networks are described in detail in DeRosa et al. (2024).

#### Analyses during rest

New to this report, we characterize the representational similarity patterns across brain regions during rest by combing data from two resting-state fMRI runs for each participant. As with the analysis during WM operations, we applied the Glasser parcellation (Glasser et al., 2016) to extract multivoxel activation patterns from each of the 360 cortical parcels. Activation patterns were standardized prior to further analysis to ensure consistency across parcels and sessions.

The combined resting-state fMRI runs were segmented into non-overlapping windows of 30 seconds. Using this 30-second window, each segment contained approximately 65 time points, given the repetition time (TR) of 0.46 seconds. Across the two resting-state runs, which totaled 11.5 minutes (approximately 5 minutes and 45 seconds per run), 23 windows were generated per subject. Within each window, Pearson correlation matrices were computed to capture the similarity of activation patterns across the 360 cortical parcels defined by the Glasser parcellation. The lower triangular portion of each correlation matrix was extracted to isolate unique pairwise relationships between parcels. This procedure was repeated for all 23 windows. Finally, Pearson correlations were calculated between the lower triangular values of the correlation matrices from different windows to quantify representational similarity across time windows, producing a RSA matrix for each subject. For additional details on why we chose a 30-second window over the 2.75-second window used in the operation RSA, see **Supplemental Materials**.

### Functional Connectivity Analysis

New to this report, we performed functional connectivity analyses using CONN (Whitfield-Gabrieli & Nieto-Castanon, 2012) (RRID:SCR_009550) release 22.a (Nieto-Castanon & Whitfield-Gabrieli, 2022) and SPM (Friston et al., 2010) (RRID:SCR_007037) release 12.7771. FC patterns for the four WM-control operations and resting-state data were characterized using ROI-to-ROI connectivity matrices from the Glasser parcellation (Glasser et al., 2016) that encompassed 360 ROIs. Connectivity strength was represented by Fisher-transformed bivariate correlation coefficients derived from a weighted general linear model (weighted-GLM) (Nieto-Castanon, 2020b). We employed Generalized FC, a method developed by Elliott et al. (2019), which generated FC maps for the four WM-control operation datasets by regressing out effects of rest and other nuisance covariates (such as motion and physiological regressors). These coefficients were calculated for each pair of seed and target areas for each of the four WM-control operations, modeling the association between their BOLD signal timeseries. Individual scans were weighted by a boxcar signal representing each WM-control operation, convolved with an SPM canonical hemodynamic response function, and rectified. The same approach was used for resting-state connectivity but without the WM-control operation weighting. Refer to the **Supplemental Materials** for the preprocessing and denoising procedures.

### Dimensionality Reduction of both RSA and Functional Connectivity Matrices

New to this report, we subjected each of the RSA and FC matrices (RSA matrix, FC matrix during WM-control operations, FC matrix during rest) to diffusion map embedding (see **Supplemental Materials**). We did so because these matrices are high-dimensional data (e.g., 360 by 360 FC matrix), and diffusion map embedding allows to minimize the influence of noise relative to signal and to enable better visualization of our results. This procedure reduces the patterns to underlying principal axes of representational variation or FC (Huntenburg et al., 2018), which have been referred as cortical gradients (Margulies et al., 2016). More specifically, the use of gradients enhances the signal-to-noise ratio by projecting high-dimensional data into a lower-dimensional space that captures the most meaningful and relevant patterns. This process reduces random trial-level variability, which would otherwise obscure meaningful neural patterns, and focuses on the primary components that represent true neural variability and organization (Hong et al., 2020; Huntenburg et al., 2018; Peraza et al., 2024; Zhang & Zang, 2023).

Each individual’s RSA, FC WM-control operation (maintain, replace, suppress, and clear), and FC resting-state 360×360 correlation matrix were thresholded to retain the top 10% of the connections. Then, the regional representational similarity, or functional connectivity profile similarities were calculated using row-wise cosine similarity for symmetry. The resulting normalized angle matrix was subjected to diffusion map embedding, a non-linear dimensionality reduction technique. In this new space, cortical nodes strongly interconnected by either many suprathreshold edges or a few very strong edges are closer together, while nodes with little or no inter-covariance are farther apart. In this context, inter-covariance extends the concept of covariance to describe the similarity between entire representational similarity or connectivity profiles of regions (e.g., rows in the matrix) rather than just pairwise relationships. While covariance measures how two variables co-vary, inter-covariance emphasizes how multi-voxel activation or functional connectivity patterns across multiple regions are related. This approach captures higher-order relationships across the brain, providing a better understanding of similarity in activation patterns or connectivity between brain regions beyond simple pairwise associations.

The first three gradients from the diffusion map embedding were selected for further analysis because they accounted for over 50% of the variance across each of the RSA matrices of the four WM-control operations, the FC matrices of the four WM-control operations, and the resting-state analyses using the Brainspace package in Python (Vos de Wael et al., 2020). This amount of variance was considered sufficient to capture the most meaningful patterns related to the neural organization of WM-control operations. These three group-level gradient components across all four operations were then aligned using joint embedding (Nenning et al., 2020). The 3D space from these gradients allowed clear visual inspection, differentiation, and further analyses of the representations and connectivity within the four _wmo_networks. Embedding solutions for each individual were aligned to the group-level hold-out embedding via joint embedding. See **Figure 2** for an illustration of the computations applied to the RSA and FC matrices to derive the gradient solutions. The same thresholding, diffusion map embedding, and alignment procedures were applied to the RSA and FC matrices to derive the three gradients.

**Figure 2.**
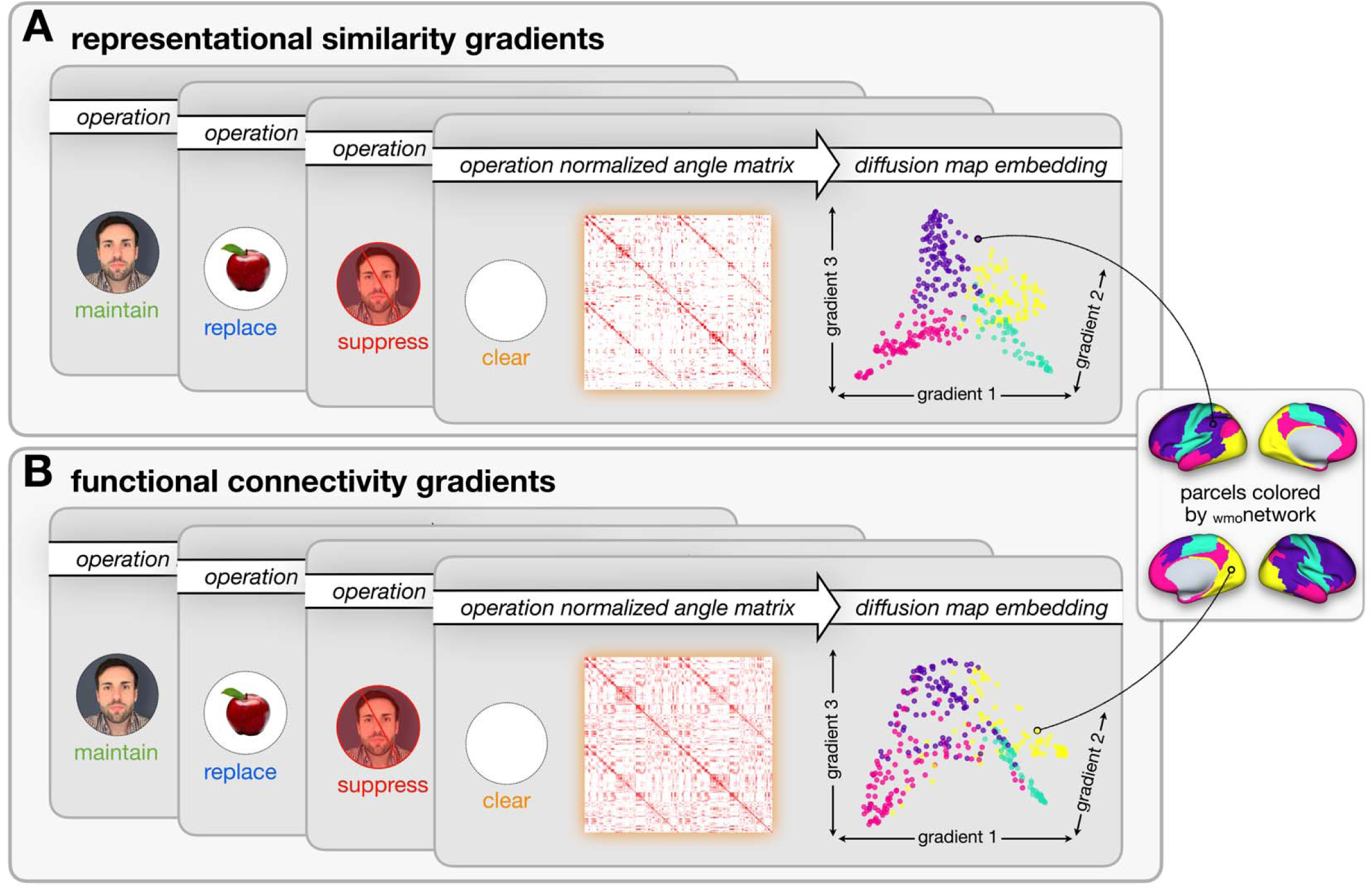
Dimensionality reduction to derive gradients of A) representational similarity and B) functional connectivity patterns across the WM-control operations: (maintain, green; replace, blue; suppress, red; clear, orange). Both A and B illustrate the gradient group-level diffusion map embedding solutions for illustration purposes. Parcels are colored by DeRosa et al. (2024) _wmo_network: yellow, V_wmo_; teal, SM_wmo_; pink, DM_wmo_; purple, FP_wmo_. To derive the individual-level gradients for each participant, the normalized angle matrix across 360 Glasser parcels wa thresholded to retain the top 10% of the connections for each of the four WM-control operations, as well resting-state (not shown in this figure). Then, the regional similarity (RSA), as well a functional connectivity profile similarities were calculated using row-wise cosine similarity for symmetry. Separately, the normalized angle matrixes were subjected to diffusion map embedding, and the top three components were extracted.

The gradients derived from RSA and FC analyses revealed three similar, but not identical, main axes. As a brief overview, gradient-1 primarily captures the differentiation between sensory-motor and higher-order associative regions, gradient-2 delineates the distinctions between visual and frontoparietal areas, and gradient-3 reflected differences between frontoparietal control regions and default mode regions. The **Supplemental Materials** provide further details on the similarities and differences between the RSA and FC gradients.

### Variability of Neural Representations and Patterns of Connectivity

To examine how individual differences in difficulty in controlling thoughts might relate to neural patterns, which is the focus of the current report, we need a method to examine variability in those neural patterns. We do so with reference to the three main gradients derived for each of our matrices (RSA matrix, task-based FC, FC matrix at rest), examining two aspects of neural variability. The first, eccentricity, examines how distinct (or overlapping) the patterns for a given _wmo_network (e.g., V_wmo_) are from the other three _wmo_networks. The second, dispersion, examines how variable the parcel patterns are within a given _wmo_network (e.g., the FPC_wmo_).

### Eccentricity

Eccentricity (Del Río et al., 2024; Valk et al., 2020) measures the distance between each parcel and the center of the gradient coordinate system across all _wmo_networks. This metric was computed individually for each participant using their first three gradients. This approach places each parcel in the three-dimensional gradient space, where the average eccentricity value for each _wmo_network reflects its segregation from other _wmo_networks. In other words, average eccentricity for a given _wmo_networks measures how distinct the neural patterns are for each _wmo_network from the rest of the brain’s neural patterns. Higher values indicate greater segregation and more differentiation, while lower values suggest more overlap between the networks.

In RSA, higher eccentricity of a _wmo_network implies greater representational segregation, suggesting that the _wmo_network represents the operation more distinctly compared to the patterns of the rest of the brain. Similarly, in functional connectivity (FC), higher eccentricity is thought to reflect greater functional segregation, indicating that a _wmo_network operates more independently by having connectivity patterns that are more distinct and less correlated with those of other _wmo_networks.

### Dispersion

Dispersion (Bethlehem et al., 2020; Cross et al., 2021; Del Río et al., 2024) captures the variability of the neural patterns within networks (as compared to eccentricity which examined patterns across networks). Within-network dispersion is the sum of the Mahalanobis distances of each region’s gradient score from the _wmo_network centroid (i.e., the median coordinates in the 3D gradient space of all parcels belonging to that _wmo_network). Mahalanobis distance was chosen over Euclidean distance for these calculations due to its ability to account for the different variances in each dimension of the gradient space, which is important for capturing the relationship between brain regions in our multi-dimensional space. Within-network dispersion is particularly useful when focusing on a single _wmo_network’s internal organization, independent of the overall pattern of the organization of other _wmo_networks.

In RSA, higher within-network dispersion indicates greater variability in how the regions within a network represent each WM-control operation. In FC, higher within-network dispersion reflects greater variability in the FC patterns across parcels within a network. **Figures 3** (eccentricity) and **4** (dispersion) show these two types of variability and their hypothesized relationships to difficulties in controlling thoughts.

**Figure 3.**
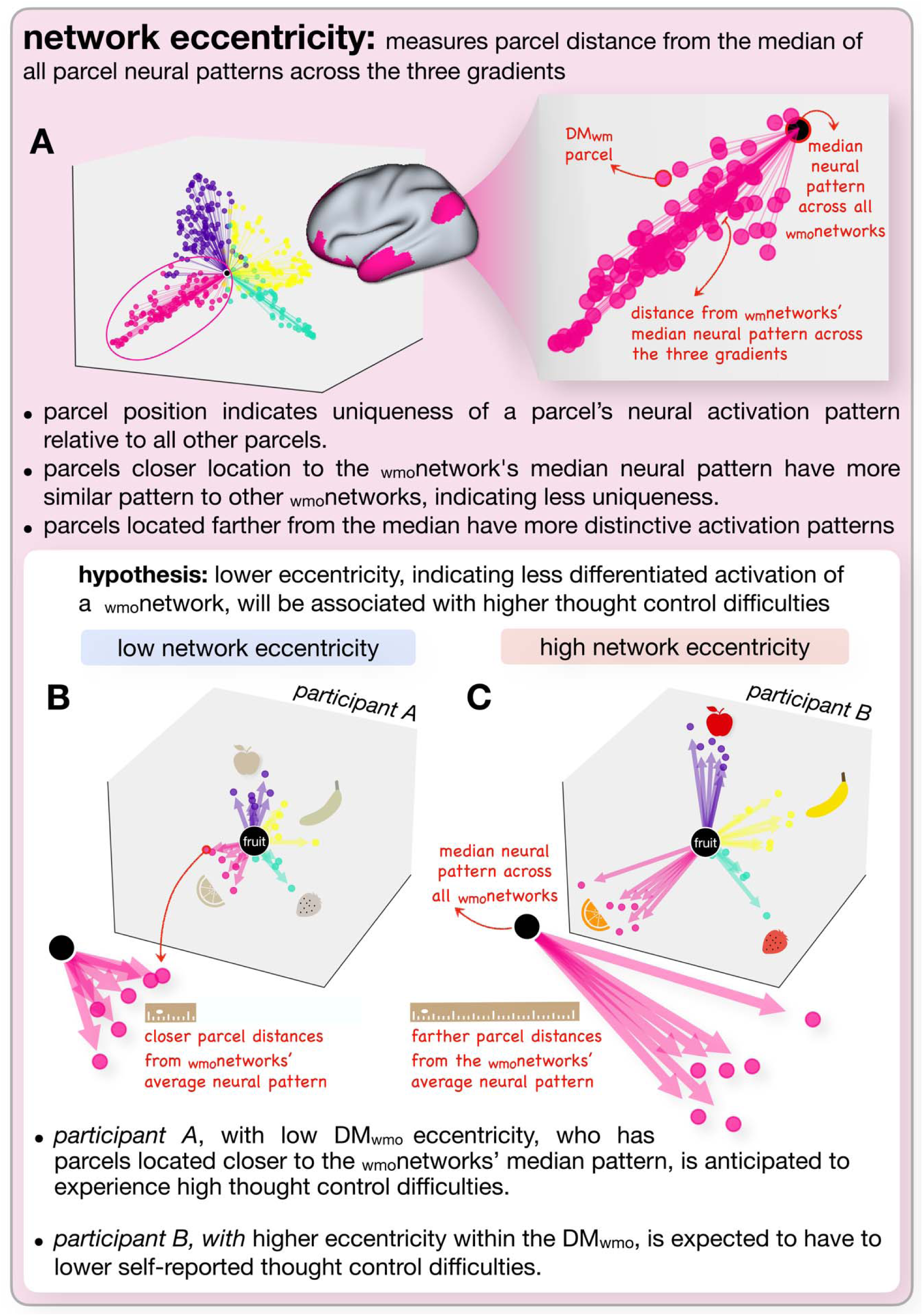
Hypothesis-driven representation of _wmo_network eccentricity. This figure illustrates the concept of _wmo_network eccentricity and its hypothesized relationship with thought control difficulties. A) (left) displays group-level metrics for a sample WM operation, showing the general organization of _wmo_networks. The _wmo_networks are color-coded as per DeRosa et al. (2024): yellow for the V_wmo_, teal for the SM_wmo_, pink for the DM_wmo_, and purple for the FP_wmo_. The magnified view on the right focuses on the DM_wmo_, with the large central dot representing the average neural pattern across all _wmo_networks. Individual DM_wmo_ parcels, shown as pink dots, are positioned according to their distance from this central average, illustrating eccentricity. B) Shows data from hypothetical Participant A, characterized by low _wmo_network eccentricity within the DM_wmo_. C) Shows data from hypothetical Participant B, characterized by high _wmo_network eccentricity within the DM_wmo_. The ruler icons illustrate distance, where a shorter ruler indicates a shorter distance from the median, while a longer ruler indicates a farther distance. The variations in the fruit across Participants A and B are shown as an example to illustrate differences in neural patterns across the _wmo_networks, highlighting the specificity of each _wmo_network in relation to the overall pattern of brain activity.

### Analysis Procedure for Identifying the Neural Correlates of Thought Control Difficulties

Once again, as an overview, we examined how individual differences in self-reported thought control difficulties are associated with distinct types of neural metrics: univariate activation, whole-brain classifier accuracy scores, and the cortical gradient-based metrics (eccentricity, dispersion) derived from _wmo_network analyses. Hierarchical multiple regression models assessed the additional explanatory power of _wmo_network organization and FC characteristics beyond the baseline accuracy of differentiating WM-control operations in predicting individual differences in thought control difficulties. False Discovery Rate (FDR) correction (Benjamini & Hochberg, 1995) was applied to all regression models for each neural metric (e.g., eccentricity, dispersion). Specifically, for each neural metric, we performed 16 separate regression models (one for each combination of the four _wmo_networks by four WM-control operations). The Spearman rank-order correlations for all WM-control and resting-state neural metrics, including WM-operation classifier accuracy and _wmo_network organization, are provided in **Supplemental Materials Figures 3-9)**.

### Univariate Brain Activation

To determine the relationship between univariate brain activation and thought control difficulties, each participant’s thought control difficulties score was included as a covariate of interest in the univariate models. All statistical inferences were carried out using permutation testing implemented via FSL’s randomise (version 2.9) with 10,000 permutations. Analyses were restricted to voxels within a gray matter mask. First, a voxel-wise threshold of *p* < 0.001 (uncorrected) was applied. Next, a cluster-forming threshold of *p* < 0.001 (uncorrected) was used. FWE correction was applied at the cluster level, and results were considered statistically significant if they survived a corrected threshold of *p* < 0.05.

### Whole-Brain Classifier Accuracy Scores for each Working Memory Operation

We implemented a series of regressions to examine whether the multi-voxel activation patterns of each WM-control operation (as determined by whole-brain classifier accuracies) varied as a function of an individual’s thought control difficulties. We conducted separate regressions for each WM-control operation’s classifier accuracy score to predict thought control difficulties. Following these analyses, we included all four classifier accuracy scores in a multiple regression model to examine the unique variance that each WM-control operation contributes to thought control difficulties when controlling for the other operations. This approach allowed us to determine whether any specific operation significantly explains thought control difficulties beyond the shared variance across all operations.

### Cortical Gradient _wmo_Network Organization

We implemented a series of regressions to investigate whether an individual’s self-reported difficulties in controlling thought varied as a function of cortical gradient (eccentricity and dispersion) organization. Separate regressions were conducted for each WM-control operation, examining the relationship between self-reported thought control difficulties and _wmo_network eccentricity as well as dispersion characteristics for the RSA metrics. Identical analyses were then performed for the connectivity gradients during the WM-control operations. Lastly, to examine the specificity of any observed effects, we used the same approach and metrics as above to determine whether similar relationships were present during the resting-state.

### Examining the Relative Power of Distinct Brain Metrics for Predicting Individual Differences with Difficulty Controlling Thoughts

Our final goal was to examine the extent to which different classes of _wmo_network gradient organization provided additional value beyond the whole-brain classifier accuracy model in predicting individual differences in the difficulty of controlling thoughts. The analyses were designed to evaluate the incremental explanatory power of these metrics by first assessing the whole-brain classifier accuracy and then progressively incorporating _wmo_network RSA gradient metrics (eccentricity and dispersion) to predict individual differences in the difficulty of controlling thoughts. Once the added value of these RSA metrics was established, the _wmo_network connectivity gradient organization was introduced to further assess its contribution to predicting thought control difficulties. Ultimately, we wished to determine the maximal amount of variation in self-reported thought control difficulties that could be accounted for by multiple metrics. To investigate these issues, we employed a hierarchical multiple regression modeling approach that systematically introduced the class of brain metrics in a stepwise manner.

### Stage 1: Model 1a (Principal Component of Whole-Brain Classifier Accuracy)

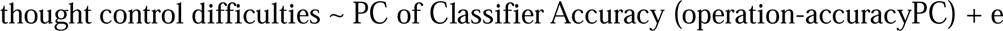

At the first level, we determined whether the degree to which the classifier accuracies for the WM operations could predict individual differences in thought control difficulties. As the classifier accuracies were highly correlated across the four operations (see **Results**), we conducted a principal component analysis (PCA) on the four WM-control operation accuracy scores to extract a single principal component (PC) that captures the shared variance in the ability to differentiate between the WM-control operations. Based on prior recommendations that the sample size should be at least five times the number of variables, our sample size was appropriate for deriving the PCA from the four measures (Hatcher, 1994).

### Stage 2: Models 1a + 1b (Adding _wmo_Network RSA Cortical Gradient Metrics)

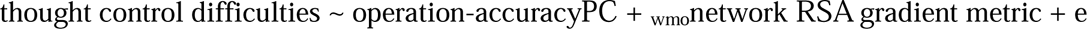

In this next stage, we assessed whether _wmo_network RSA metrics (eccentricity and dispersion) provided additional explanatory power for predicting thought control difficulties beyond the baseline model that included the PC of the classifier accuracy (operation-accuracyPC). As described above, we conducted separate regression analyses for each RSA gradient metric (16 for dispersion (four _wmo_network by four operations) and 16 for eccentricity) to identify metrics that significantly predicted thought control difficulties. From those regressions, the _wmo_network RSA gradient metrics (e.g., eccentricity or dispersion for a given operation) that were significantly associated with thought control difficulties (*p_FDR_* < .05) were carried forward for the current nested multiple regression analyses.

For the Stage 2 analyses, each significant RSA gradient metric was included as an additional predictor in a nested multiple regression model alongside the PC, resulting in separate models for each metric. Thus, for this stage, each nested model predicted thought control difficulties using the PC and one significant _wmo_network RSA gradient metric. Seven such nested models were constructed, as seven RSA metrics were found to be significant in the initial regressions (see **Results**).

To evaluate whether adding _wmo_network RSA gradient metrics improved model fit (i.e., the ability of the model to explain variance in thought control difficulties), we compared the nested models (PC + RSA metric) to the baseline PC-only model using the F-test for nested models. A *p* < 0.05 from the F-test indicated that the nested model with these additional neural metrics explained significantly more variance in thought control difficulties compared to the simpler model, which, in this case, was the PC-only model. Improvements in adjusted R^2^ were used to ensure that the RSA metrics explained additional variance without artificially inflating it. Nested model comparisons (e.g., 1a vs. 1a + 1b) explicitly confirmed whether RSA gradient metrics added significant explanatory power.

### Stage 3: Models 1a + 1b + 2b (Adding _wmo_Network Connectivity Cortical Gradient Metrics)

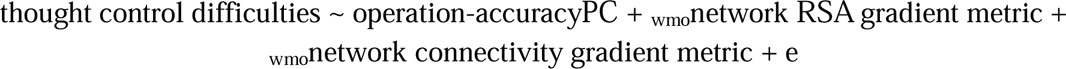

In the final stage, a single _wmo_network connectivity gradient metric was added to the significantly better-nested models from Stage 2 to evaluate whether this connectivity metric provided additional explanatory power beyond the combined PC and RSA gradient metrics. This connectivity metric was included because it was the only one significantly associated with thought control difficulties (*p_FDR_* < .05) in the initial regressions (see **Results**). Adding this metric to the respective significant Stage 2 models resulted in nested models with three predictors: PC of the classifier accuracy, a _wmo_network RSA gradient metric, and the _wmo_network connectivity metric.

Nested model comparisons were conducted specifically between the significant Stage 2 models (PC + RSA gradient metric) and their corresponding Stage 3 models (PC + RSA gradient metric + connectivity gradient metric) using the F-test for nested models (e.g., 1a + 1b vs. 1a + 1b + 2b). Adjusted R^2^ values were used to assess whether adding the connectivity gradient metric explained additional variance in thought control difficulties beyond what was captured by the Stage 2 models.

Lastly, to determine whether the effects noted in the models above are specific to brain metrics while individuals are engaged in WM operations, we also conducted regression analyses to determine whether resting-state metrics (dispersion and eccentricity) provide additional predictive power for thought control problems beyond task-based measures. In each analysis, we added resting-state metrics (matched by _wmo_network and operation) to the task-based models and used nested F-tests to assess changes in explanatory variance. Detailed methods and findings are presented in the **Supplemental Materials** because these resting-state metrics did not add additional explanatory power.

## Results

### Univariate Activation During the WM-Control Operations

The relationship between an individual’s level of thought control difficulties and univariate activation revealed just two significant clusters in BA 24, one of the contrast of maintain vs. clear and the other for the contrast of clear vs. the other WM operations (see **Supplemental Materials** for further details). These limited findings indicate that our consideration of multi-variate methods to explore brain-behavior associations are merited.

### Whole-Brain Classifier Accuracy Scores of Working Memory Operations

Individuals who reported higher levels of difficulty in controlling their thoughts exhibited a lower classification accuracy score for a given operation (vs. the other three operations) for each of the maintain, suppress, and clear operations. A similar but non-significant relationship was observed for the replace operation (*p_FDR_* > .1) (see **Figure 5** and top section **Supplemental Table 1**).

**Figure 4.**
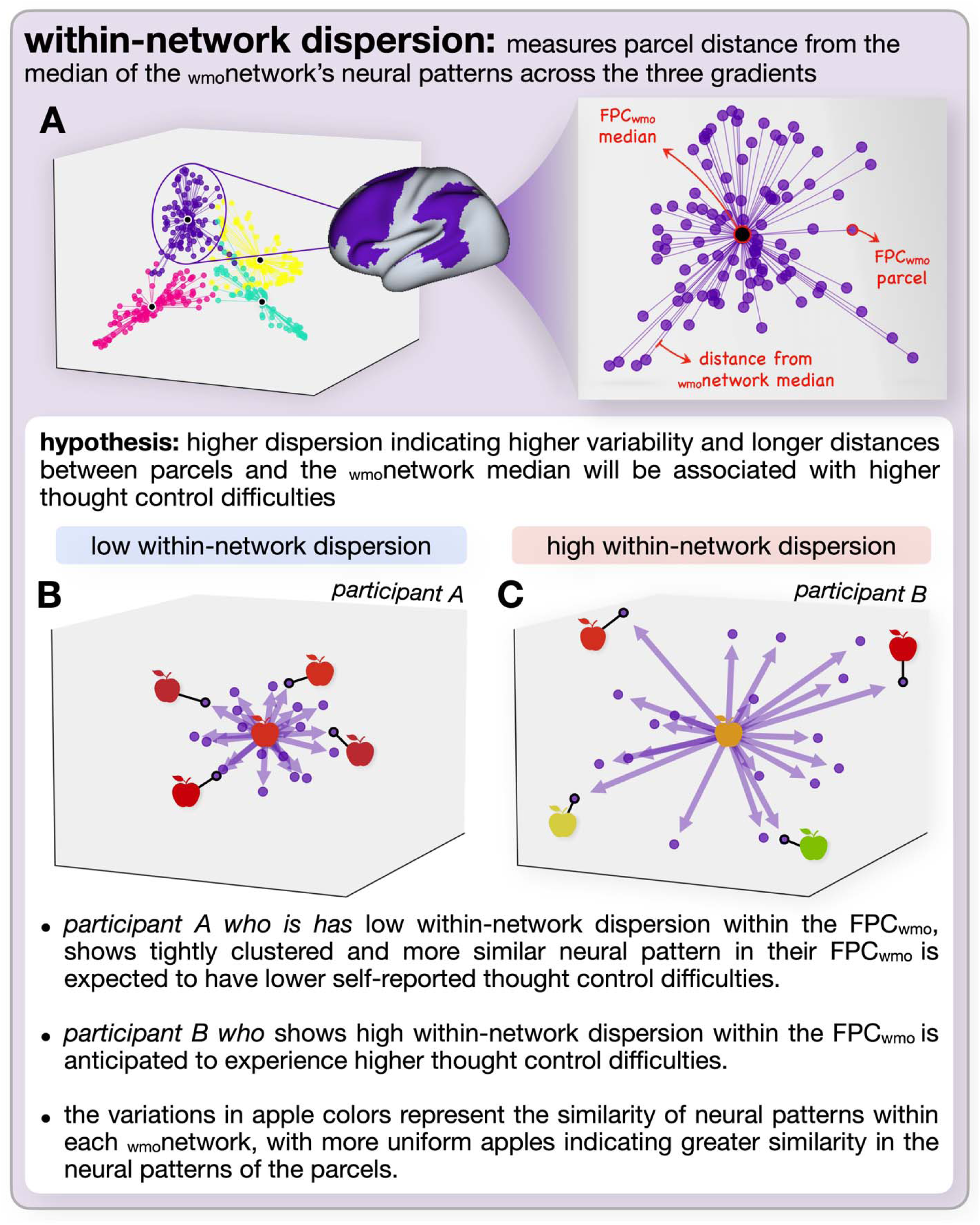
Hypothesis-driven representation of within-network dispersion. This figure illustrates the concept of within-network dispersion and its hypothesized relationship with thought control difficulties. A) (left) presents group-level metrics for a sample working memory operation, showing an overall view of _wmo_network organization. _Wmo_networks are color-coded according to DeRosa et al. (2024): yellow for the V_wmo_, teal for the SM_wmo_, pink for the DM_wmo_, and purple for the FP_wmo_. The enlarged view on the right highlights the FPC_wmo_, with the large central dot representing the median pattern across FPC_wmo_ parcels and the smaller purple dots representing individual FPC_wmo_ parcels. The lines connecting each parcel to the median indicate the Mahalanobis distance across three gradients, reflecting dispersion within the _wmo_network. B) Shows data from Participant A, characterized by low within-network dispersion within the FPC_wmo_. C) Shows data from Participant B, characterized by high within-network dispersion within the FPC_wmo_. The variations in apple shapes and colors represent the degree of similarity among neural patterns within each _wmo_network, with more uniform apples indicating greater similarity in parcel patterns.

**Figure 5.**
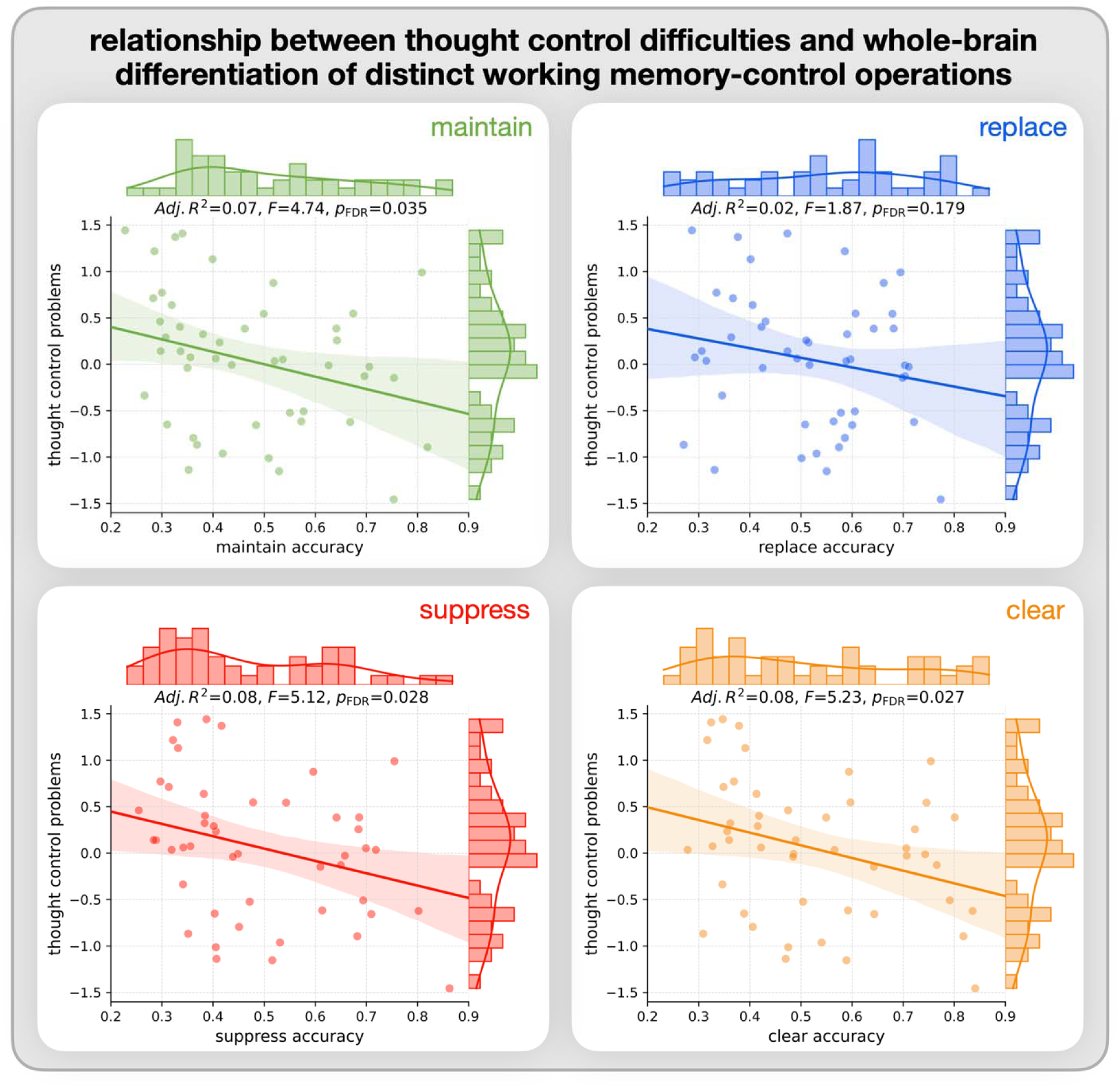
Regression relationships between working memory operation accuracy and thought control difficulties. Each plot represents one working memory operation: maintain (green, top left), replace (blue, top right), suppress (red, bottom left), and clear (orange, bottom right). Points indicate individual participant data, with thought control problem scores plotted against the accuracy score of each operation. The regression line represents the linear association, with shaded areas indicating the confidence intervals. Regression statistics (Adj. R², F-value, and FDR corrected p-value) are provided on the top of each subplot.

### RSA Cortical Gradient Measures of _wmo_Network Organization

With regards to network eccentricity, higher thought control difficulties were associated with lower DM_wmo_ eccentricity and higher V_wmo_ eccentricity during the suppress operation. With regards to within-network dispersion, higher thought control difficulties were associated higher within-network dispersion within the FPC_wmo_ during the suppress operation and within higher within-network dispersion of the DM_wmo_ during the maintain, replace, and clear operations (**Table 1**). Scatter plots for DM_wmo_ eccentricity during the suppress condition and FPC_wmo_ within network dispersion during the suppress operation are provided in Figure 6. In general, these findings indicate that the less distinct the neural representations within a specific _wmo_network (e.g., DM_wmo_) are from the representations in other _wmo_networks (as indexed by eccentricity), and the more variable these representations are within the _wmo_network itself (as indexed by dispersion), the more difficulty an individual reports in controlling their thoughts. See the middle section of **Supplemental Table 1** for a complete report of all regression results.

**Figure 6.**
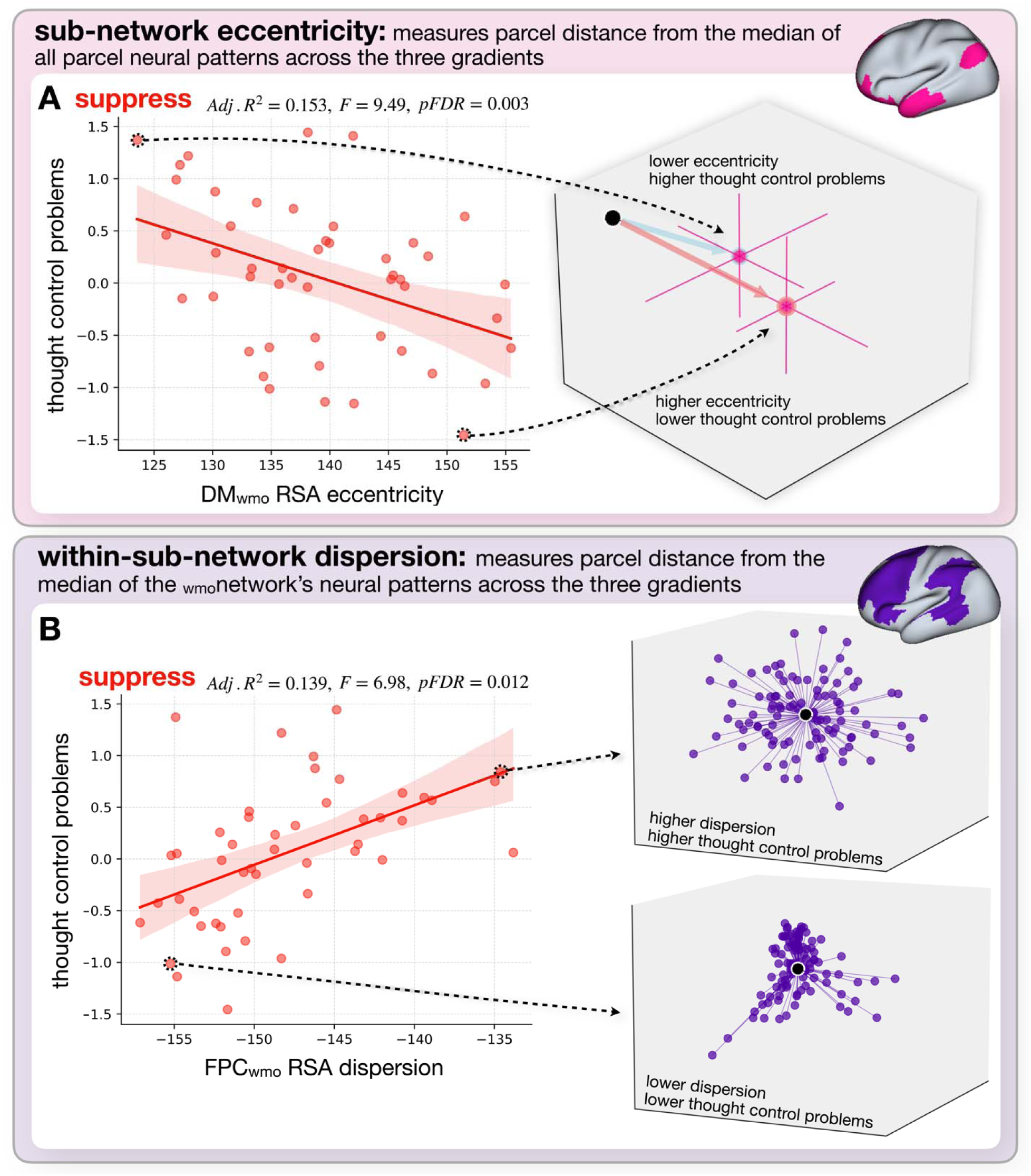
Relationship between RSA _wmo_network eccentricity and within-network dispersion with thought control difficulties. This figure illustrates the regression relationships between RSA metrics (eccentricity and within-network dispersion) for the suppress operation and thought control difficulties, showing _wmo_network neural pattern variations among participants. A) and B) (left) display regression plots for DM_wmo_ RSA eccentricity (Panel A) and FPC_wmo_ RSA within-network dispersion (Panel B) against thought control problem scores. In each plot, points represent individual participant data, with thought control difficulties plotted against RSA metric values. Linear regression lines indicate the association, with shaded areas denoting confidence intervals, and regression statistics (Adj. R², F-value, and p-value) are shown on the top of each subplot. The right side of each panel provides illustrative examples of neural pattern metrics for two selected participants. In A) DM_wmo_ eccentricity is shown for one participant with high eccentricity (red arrow, lower thought control problems) and one with low eccentricity (blue arrow, higher thought control problems). Pink dots represent mean eccentricity values across three gradients, pink lines show standard error bars for parcel positions, and the black dot indicates the _wmo_network’s average neural pattern. In B) FPC_wmo_ dispersion is illustrated for a participant with high dispersion, shown by a broader scatter of purple dots (parcels) in the FPC_wmo_, and a participant with low dispersion, indicated by a tighter clustering of parcels around the FPC_wmo_ median.

**Table 1.** presents the significant regression outputs showing the relationships between thought control difficulties and who brain WM-control operation classifier accuracy, _wmo_network RSA organizational patterns, and _wmo_network functional organizational patterns. The p-values in this table reflect the statistical significance of the relationship marked by asterisks(\**p_FDR_*< 0.05, ** *p_FDR_* < 0.01) indicating significant associations.

| Data | Operation | Metric | Network | $F$ | Adj. $R^2$ | $t$ | $p_{FDR}$ |
| --- | --- | --- | --- | --- | --- | --- | --- |
| whole-brain classifier | maintain | accuracy |  | 4.737 | 0.074 | -2.176 | 0.035 |
|  | suppress |  |  | 5.115 | 0.081 | -2.262 | 0.028 |
|  | clear |  |  | 5.229 | 0.083 | -2.287 | 0.027 |
|  | operation-accuracyPC |  |  | 4.99 | 0.078 | 2.234 | 0.03 |
| representational similarity | maintain | dispersion | $DM_{wmo}$ | 6.921 | 0.112 | 2.631 | 0.012 |
| | replace | dispersion | $SM_{wmo}$ | 4.972 | 0.097 | -2.23 | 0.032 |
| | replace | dispersion | $DM_{wmo}$ | 4.347 | 0.066 | 2.085 | 0.043 |
| | suppress | eccentricity | $DM_{wmo}$ | 9.494 | 0.153 | -3.081 | 0.003 |
| | suppress | dispersion | $FPC_{wmo}$ | 6.975 | 0.139 | 2.641 | 0.012 |
| | clear | dispersion | $DM_{wmo}$ | 4.404 | 0.084 | 2.099 | 0.043 |
| connectivity | | eccentricity | $DM_{wmo}$ | 12.34 | 0.235 | -3.513 | 0.001 |

### Connectivity Cortical Gradient Measures of _wmo_Network Organization

These analyses only yielded one significant result, which was that greater thought control difficulties were associated with DM_wmo_ connectivity patterns that were less differentiated from other _wmo_networks (i.e., lower eccentricity) during the maintain operation (**Table 1**). See the bottom section of **Supplemental Table 1** for a complete report of all connectivity regression results.

### Specificity of _wmo_network Cortical Gradient Organization to Control over Working Memory

None of the resting-state _wmo_network RSA or connectivity gradient organization metrics revealed significant associations with thought control difficulties. These findings suggest that the _wmo_network neural organization related to thought control difficulties is more likely linked to an individual’s ability to control information in WM rather than reflecting a general characteristic of overall brain (dis)organization (**Supplemental Table 2**).

### Hierarchical Multiple Regression Models

#### Stage 1: Principal Component of the Whole-Brain Classifier Accuracy

As discussed in the methods section, because the operation accuracy scores were highly correlated, we created a PC (operation-accuracyPC) from the four WM-control accuracy scores, which explained 88% of the variance across operations and significantly predicted thought control difficulties (*F*=4.99, Adj. R^2^=.078; Table 1). This model indicated that the PC accounted for approximately 8% of the variance in thought control difficulties, providing a benchmark for evaluating the impact of adding the representational and connectivity gradient metrics.

#### Stage 2: Adding _wmo_Network Representational Similarity Gradient Metrics)

Stage 2 examined whether adding the significant characteristics of the _wmo_network RSA gradient neural representations to the baseline operation-accuracyPC model would improve the variance in explaining thought control difficulties. Three of the seven nested models significantly explained more variance in thought control difficulties than the PC alone (**Supplemental Tables 3-4**). Of these three significant models, two included information about the pattern of the representations from the DM_wmo_ (suppress eccentricity and maintain within-network dispersion) and one about the pattern of representations from the V_wmo_ (suppress eccentricity). All three significant models—operation-accuracyPC + DM_wmo_ suppress eccentricity, operation-accuracyPC + DM_wmo_ maintain within-network dispersion and operation-accuracyPC + V_wmo_ suppress eccentricity—each independently improved over the PC model alone, explaining approximately 14%, 20%, and 15% of the variance in thought control difficulties, respectively. To determine whether the DM_wmo_ maintain dispersion and DM_wmo_ suppress eccentricity measures account for unique variance in predicting thought problems, we conducted variance inflation factor (VIF) analyses by including them in the same models and comparing their contributions for predicting thought control problems. These analyses revealed low multicollinearity (VIF ≈ 1), indicating that each measure contributes distinct variance for predicting thought control difficulties.

#### Stage 3: Adding _wmo_Network Connectivity Gradient Metrics)

The final stage investigated whether adding the only significant _wmo_network connectivity gradient pattern—eccentricity of the DM_wmo_ during maintain—could further enhance the ability to explain thought control difficulties. Adding DM_wmo_ eccentricity to the three significant models from Stage 2 further improved the explanatory power of all three models tested. These models explained approximately 27%, 29%, and 31% of the variance in thought control difficulties, respectively. Hence, combining neural representations and connectivity patterns best explains individual differences in thought control (**Supplemental Tables 3-4).**

In sum, combining the _wmo_network RSA gradient neural representational characteristics with the _wmo_network connectivity gradient patterns significantly improved the ability to explain individual differences in thought control difficulties above and beyond the operation classifier PC. Two consistent findings emerged specifically across the hierarchical regression models: suppression-related _wmo_network organization—particularly within the DM_wmo_—was a strong predictor in understanding individual differences in thought control difficulties, and the predictive effect of the PC was attenuated when the _wmo_network organization metrics were included (**Supplemental table 4**). Among the planned hierarchical regression models, the one that explained the most variance in self-reported thought control difficulties, accounting for approximately 30% of the variance, included both RSA gradient and connectivity gradient metrics of _wmo_network organization. Finally, an exploratory multiple regression model that included the PC and all eight of the significant _wmo_network organization metrics explained about 50% of the variance in thought control difficulties, outperforming all other models in nested comparisons (**Supplemental Table 4**). These findings suggest that the various multivariate metrics detect distinct neural processes associated with difficulties in controlling thoughts and can account for a fair amount of that variation.

## Discussion

The main result of the current study demonstrates that individual differences in self-reported difficulties in controlling thought are associated with differences in the multivariate neural patterns of activation that underlie how the brain implements different WM-control operations (e.g., maintain, suppress). This is a novel finding. In particular, our analyses showed that less defined and less consistent representational structures across networks during these WM-control operations are linked to greater difficulties in controlling thought. We demonstrated this relationship in three distinct manners. First, our results indicated that the poorer an individual’s ability to control their thoughts, the less distinct each of the WM operations was from the other three as indexed by the accuracy of a whole-brain machine learning classifier for each operation. Second, we found that one’s self-reported ability to control thoughts is associated with how the WM-control operations are represented in specific _wmo_networks. These effects were observed mostly for the DM_wmo_, although effects were also observed for the V_wmo_ and FPC_wmo_, and mostly with the suppress operation. Finally, greater difficulties in controlling thought were associated with one aspect of the connectivity pattern with the DM_wmo_. Importantly, these associations were observed explicitly for the task (i.e., when WM-control operations are employed) but not with resting-state. This finding suggests that difficulties in thought control are primarily related to adaptations in the organization of neural activity during the specific control of information in WM rather than being due to general disorganization in brain connectivity as gleaned from resting-state data. The remainder of this discussion will consider the specific results in more detail, explore their implications, and place them within the framework of existing literature.

Consistent with our hypothesis, individuals reporting higher difficulties in controlling their thoughts exhibited less distinct neural activation patterns as indicated by classifier accuracy across three of the WM-control operations — maintain, clear, and suppress. In other words, the more consistently someone can implement distinct forms of control over the information in their WM, the better they are at managing their thoughts. While the classifier accuracy for the replace operation showed the same trend, the effect was not significant. Upon closer inspection, we found that the distribution of these accuracy scores was more uniform across participants (see distribution on the top of the replace operation scatter plot in **Figure 5**), and compared to the other three WM-control operations there were fewer lower classification accuracy scores, suggesting that fewer individuals struggled to differentiate replace from the other operations. Kim et al. (2020) found that the replace operation, in which an individual replaces the current item with an image previously seen, lead to quicker deactivation of the original item’s neural representation than suppressing or clearing it, providing the possibility that replace operation is not as cognitively demanding as suppress or clear.

Converging evidence for this idea was provided by the eccentricity analyses of the _wmo_networks. Eccentricity provides a metric of how different the average representation of an operation within a _wmo_network is from the average representation across all four _wmo_networks. It was observed that greater thought difficulties were associated with a greater difference (i.e., eccentricity) of the V_wmo_ from the average and less eccentricity of the DM_wmo_ during the suppress condition. This is a novel finding since, unlike most studies that have reported on BOLD activation (Chou et al., 2023) or magnitude differences in the connectivity patterns in individuals with thought-related difficulties (Burr et al., 2020; Guo et al., 2014; Hamilton et al., 2015), our analyses show that distinct configurations of distributed patterns of brain activity within specific regions are associated with individual differences in the ability to control thoughts. As to the particulars of the patterns observed, prior research (Kim et al., 2020; Brenton et al., 2024) suggests that the suppress condition may initially require selection and focused attention on the item to be suppressed, and only afterward, its actual suppression. This may help explain the pattern observed DM_wmo_: the more uniquely the DM_wmo_ represents suppression relative to other _wmo_networks, the better an individual can control the internal representation of information in WM. In contrast, the more the regions involved in visual processing differ from the brain’s average representation for suppression, the greater the difficulty with thought control. This pattern suggests that successful suppression involves disengaging from the item’s visual representation, resulting in a V_wmo_ pattern that is less distinct and more similar to the brain’s average representation during suppression.

In addition, we observed that the consistency of the representation within a _wmo_network, as measured by dispersion, is also associated with difficulties in controlling thought, with greater thought difficulties associated with greater dispersion (i.e. less consistent representations). More specifically, greater thought difficulties were associated with greater dispersion for the maintain, replace and clear operations within the DM_wmo_ and greater dispersion during the suppress operation within the FPC_wmo._ This pattern of results represents an interesting dichotomy between networks as well as operations. In our prior work (DeRosa et al., 2024), we found that only the FPC_wmo_ distinctly represented the four operations, leading us to suggest that this network may be centrally involved in the implementation of the WM operations. In addition, our prior work (Kim et al., 2020) found that only the suppress condition appears to remove information from WM (as evidenced by a lack of proactive interference on trials following a suppress operation) compared to either replace or clear. Together, these findings provide insight into why the variability of representation of the suppress operation across different parcels within the FPC_wmo_ is predictive of thought difficulties. It may be that this combination indexes the network most involved in controlling these WM operations as well as the operation that is most effective at removing information from WM. This idea is further supported by findings showing that higher cognitive control demands require more structured neural representations, particularly in frontoparietal regions (Freund et al., 2021).

The aspects of the organization of the DM_wmo_ are also involved. Within this _wmo_network, there is a dichotomy between thought control difficulties being associated with reduced dispersion during the suppress condition but increased dispersion during the other three. There is a well-documented role of the DM_wmo_ regions in internally focused cognition (Andrews-Hanna, 2012; Andrews-Hanna et al., 2014; Buckner, 2013; Menon, 2023). Our results suggest that the organization of these regions may influence the ability to manage internally focused cognition. Consequently, individuals better or worse at implementing control over thoughts may differ in how well the DM_wmo_ organizes information held in WM. Since the DM_wmo_ integrates abstract, high-level information from lower-order sensory and cognitive systems (Margulies et al., 2016; Sormaz et al., 2018), this _wmo_network may act as a hub that organizes the information being held in mind. Here, we speculate that when the representation of the operations across the different constituent pieces (i.e., parcels) of the DM_wmo_ is variable (i.e., more disperse), as is the case for the clear, maintain, and replace conditions, then a consequence is that internal representations of information are not uniform and thus more poorly controlled.

In contrast, when the representation of the suppress operation is consistent across the different constituent pieces of the DM_wmo,_ this may reflect that the suppress operation is ineffective and information remains in WM. Said differently, we speculate that while the representation of information across the FPC_wmo_ may be linked to the WM operation itself, the DM_wmo_ may represent the modulation of the internal representation of information during a given operation. Prior previous has suggested that the suppress operation may modulate the item-specific representation in WM and ultimately lead to forgetting in long-term memory (Kim et al., 2020; Bretton et al., 2024). This speculation raises important questions for future investigations: understanding why and how the stability and cohesiveness of DM_wmo_ representations influence the success of different control operations.

Finally, we observed using functional connectivity analysis that individuals with greater difficulties in controlling their thoughts showed decreased eccentricity of the DM_wmo_ during the maintain operation. This indicated a reduced distinction in connectivity patterns within this _wmo_network from the connectivity patterns throughout the rest of the brain. This finding suggests that individuals may better manage internal representations in the focus of their attention when regions within the DM_wmo_ communicate more specifically with each other rather than with regions outside this network. This increased functional connectivity may facilitate more efficient control over these representations. This finding also aligns with recent research demonstrating that brain regions within the DM_wmo_ contribute to maintaining concentration and mental engagement during cognitive tasks (Sormaz et al., 2018). Concerning the current work, the findings from Somaz et al. suggest that our DM_wmo_ regions support attention to information held in WM, which may enable better control over the mental representation of the information in focus. This proposed role of the DM_wmo_ regions in attentional support for WM content may explain why the distinctness of the configuration of this _wmo_network during the active maintenance of information in WM was related to an individual’s ability to better control their thoughts.

Importantly, it appears that these brain-behavior links are specific to brain activation during WM operations and not a more general aspect of brain organization in individuals who have difficulties in controlling their thoughts. Neither the representations nor connectivity patterns from resting-state data significantly predicted thought control difficulties. These are important and novel findings, as they suggest characteristics of the neural mechanisms supporting thought control uniquely predict thought control difficulties rather than reflecting a general state of brain disorganization.

## Limitations and Future Directions

While the current findings are novel and provocative, they are not without limitations. First, participants in this study only viewed emotionally neutral stimuli, meaning our findings are currently limited to understanding these neural representations and patterns of connectivity in the context of control over non-emotional information in WM. Since intrusive and unwanted thought patterns are typically characterized by negatively charged content (Engen & Anderson, 2018; Höping & de Jong-Meyer, 2003), the next step would be to apply our analysis framework to control negatively valenced information in WM. Second, these findings were observed in a non-clinical sample, and the degree to which they reflect brain organization during WM operations for individuals who have more severe difficulties in controlling their thoughts, especially those experienced when people meet clinical criteria for various forms of psychopathology (e.g., depression, anxiety, obsessive-compulsive disorder, substance use disorder) remains to be seen. These are issues we are working on addressing in ongoing studies.

Nonetheless, the current work has important implications for future studies aiming to improve cognitive control and reduce maladaptive thought patterns through cognitive-behavioral therapy (Moody et al., 2017; Poli et al., 2022). For example, at present, the neural consequences of CBT on patients with psychiatric disorders remain unclear (Yuan et al., 2022). Our findings provide a promising direction for understanding the neural implications of CBT and other behavioral interventions in at least two ways. First, multi-voxel pattern classifier accuracy scores that differentiate WM-control operations could serve as a reliable metric for assessing how CBT influences an individual’s ability to better differentiate between the various strategies for managing and removing thoughts from the focus of their attention. Second, examining how CBT alters the organization of neural patterns within the _wmo_networks that underlie WM thought control could help us understand how CBT modulates brain function. One way to investigate this question would be to conduct pre-and post-intervention analyses of the neural patterns within _wmo_networks, such as the DM_wmo_, involved in implementing different control operations. These analyses would provide researchers and clinicians with a quantifiable and visually interpretable method to evaluate whether interventions are improving the necessary organizational characteristics of the neural patterns for controlling or removing information from WM. Additionally, these organizational characteristics could serve as a neuro-marker for tracking progress and improvements throughout the course of a treatment intervention aimed at enhancing thought control. This trackable neuro-marker may then be able to inform more personalized and effective treatment protocols that align better with patients’ unique neural profiles and therapeutic needs (Huibers et al., 2021).

In addition, these neural markers provide potential metrics to be used in biofeedback. For example, we have found that the operation classifiers generalize across individuals within a sample (Kim et al., 2020) and across samples (Bretton et al., 2024). Hence, the metrics we have identified in the current study as associated with thought control difficulties may provide a suite of measures that can be used in real-time biofeedback studies in the magnet, where the goal of the participant is to have their pattern of brain activity match that which is associated with good control over thoughts. We are currently pursuing such research.

## Conclusions

This study is the first to demonstrate that individual differences in the ability to control one’s thoughts are specifically related to the organizational characteristics of how the brain represents control over the contents of WM. We present novel findings that advance our understanding of the neural mechanisms underlying individual differences in the ability to control thoughts. First, we show that individuals with greater difficulties in controlling their thoughts have less differentiable brain activation patterns for four distinct forms of control over information in WM. Second, we show that specific patterns of neural organization within different _wmo_networks during control over WM content are related to an individual’s self-reported difficulty in controlling their thoughts. We demonstrate that greater difficulty in controlling thought is associated with less concordance in neural representations and connectivity patterns of the parcels within the _wmo_networks across the four WM-control operations. Notably, the relationships observed regarding one’s ability to control thoughts are task-specific, as the variability and distinctiveness in neural representations are found with the WM-control operations but not during resting-state activity. This is a critical finding, as it may indicate that disruptions in thought control are more reflective of how the brain organizes information during active control over information rather than general disorganization during other mental states. We also demonstrate the ability of task-based fMRI to identify and understand individual differences, countering recent criticisms of its reliability (Elliott et al., 2020). Finally, our findings also point to potential applications for understanding the underlying neural mechanisms of psychopathological patterns, such as repetitive negative thinking, rumination, and worry. The task-specific nature of these disruptions could suggest that maladaptive thought patterns may arise from impaired neural processes during active attempts at cognitive control, making interventions that investigate these mechanisms a promising avenue for future research. Overall, this work provides a novel pathway for future studies to investigate how neural organization during control over one’s thoughts can help explain individual differences in thought control, offering promising directions for interventions to improve thought control in clinical populations.

## CRediT authorship contribution statement

**Jacob DeRosa:** Conceptualization, Methodology, Formal analysis, Investigation, Writing - original draft. **Harry Smolker:** Task design and development, Writing - review & editing. **Hyojeong Kim:** Formal analysis, Writing - review & editing. **Jarrod Lewis-Peacock:** Writing - review & editing. **Marie T. Banich:** Conceptualization, Supervision, Writing - original draft, revisions

## Data and code availability

The data are available upon reasonable request, and custom Python and bash code for all primary statistical analyses are available at: https://github.com/jakederosa123/individual_differences_thought_control

## Acknowledgments

This research was supported by R56 MH125642 and R01 MH129042 to M. Banich and J. Lewis-Peacock, MPIs.

## Declaration of Interest

Declarations of interest: none

## Notes

### Competing Interest Statement

The authors have declared no competing interest.

