## Supplemental Materials for "Multivariate Neural Markers of Individual Differences in Thought Control Difficulties"

DeRosa et al.

*Thought Control Difficulties*

For the confirmatory factor analysis (CFA), 36 ordinal variables (16 questions from the PSWQ, 5 questions from the RRS brooding subscale, 15 questions from the WBSI) regressed on age and gender were used as indicators. The CFA consisted of a bi-factor model with three questionnaire-specific factors and one common factor. Correlations between latent factors were fixed to zero, and a DWLS estimator was used. All individuals included in the model (N=1007) had complete data. This bi-factor model fit the survey data well (X2 (558)=1510, CFI=0.99, RMSEA=0.041). The bi-factor model was estimated using the cfa function, and the common factor was extracted using the lavPredict function from lavaan (version 0.6-16).

In addition, to further determine if the Penn State Worry Questionnaire (PSWQ Total), Ruminative Response Scale Brooding Subscale (RRS Brooding), and White Bear Suppression Inventory (WBSI Total) could be appropriately combined into an average score, we computed the covariances between the three thought control measures. The results show consistent variances and moderate positive covariances across the three measures, indicating that higher levels of worry, brooding, and thought suppression difficulties co-occur, supporting an average score representing general thought control difficulties.

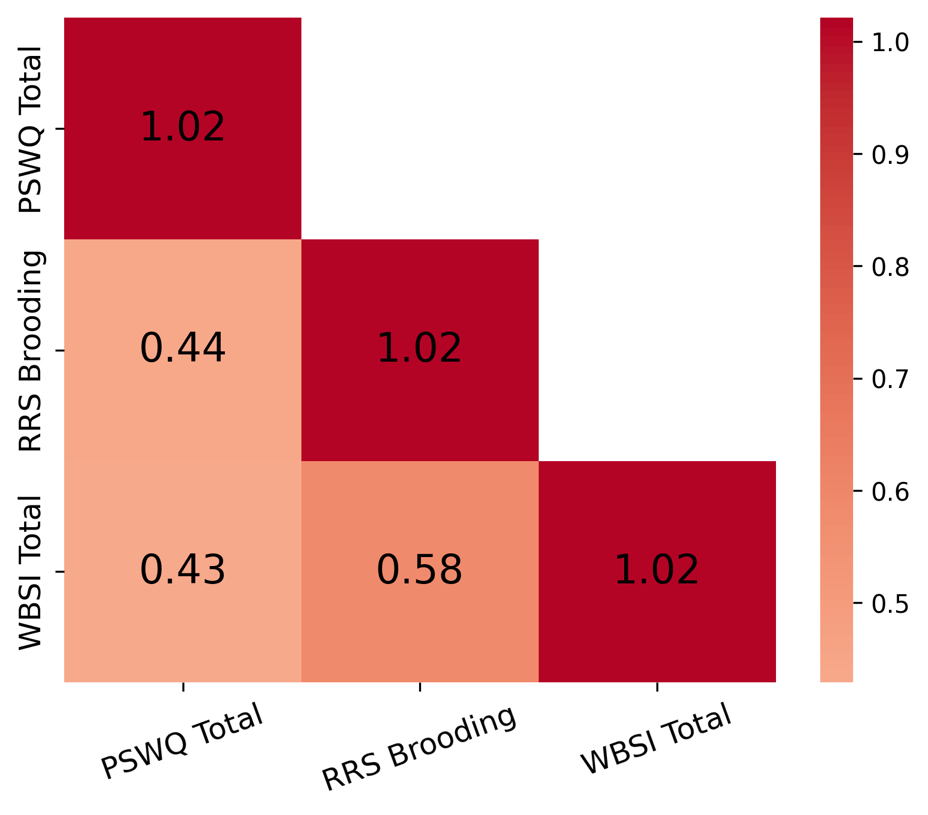

*Univariate fMRI analyses*

Our univariate activation analyses that incorporated thought control difficulties as a covariate identified two significant clusters where brain activation was associated with variations in thought control difficulties (see table below). For the contrast between Maintain and Clear, a significant cluster was observed in the left cingulate gyrus (BA24), t = 4.43, such that greater thought control difficulties were associated with increased activation in this region. For the contrast comparing the Clear condition to the other three WM operations, a significant cluster was observed in the right cingulate gyrus (BA24), t = -4.47, indicating that greater thought control difficulties were associated with reduced activation in these regions.

| Contrast | Region | BA | Max t | Voxel | X | Y | Z |
| --- | --- | --- | --- | --- | --- | --- | --- |
| Maintain vs Clear | Cingulate Gyrus (L) | BA24 | 4.43 | 434 | -2 | 20 | 30 |
| Clear All WM | Cingulate Gyrus (R) | BA24 | -4.47 | 273 | 2 | -16 | 42 |

*Resting-State Representational Similarity Analysis*

Initially, we considered using a 2.75-second window for the resting-state analysis to align with the operation-based RSA methodology. However, preliminary analyses revealed that shorter windows produced noisier and less stable correlation matrices. Stability in this context was defined as the similarity of the parcel organization derived from resting-state RSA to the parcel organization observed in task-based RSA. Specifically, we used gradient scores obtained from task-based diffusion map embeddings (discussed in detail below) as a benchmark and assessed the stability of resting-state embeddings by correlating their gradient scores with the task-based gradients. Shorter windows exhibited lower correlations with the task-based gradients, indicating reduced stability and suggesting that a 2.75-second window was insufficient for reliable representational similarity patterns in the resting-state data.

To refine the selection, we tested various window lengths ranging from 2.75 seconds to 45 seconds. The 30-second window achieved the best balance, providing the highest correlations with task-based gradient scores and producing parcel organizations most similar to those observed in task-based embeddings. Shorter windows resulted in noisier patterns with lower gradient correlations, while longer windows (e.g., 45 seconds) did not yield significant improvements in stability over the 30-second window.

*Functional Connectivity Preprocessing*

Functional and anatomical data were preprocessed using a flexible preprocessing pipeline (Nieto-Castanon, 2020a), which included realignment with correction of susceptibility distortion interactions, slice timing correction, outlier detection, direct segmentation, and MNI-space normalization, and smoothing. Functional data were realigned using SPM realign & unwarp procedure (Andersson et al., 2001), where all scans were coregistered to a reference image (first scan of the first session) using a least squares approach and a 6 parameter (rigid body) transformation (Friston et al., 1995), and resampled using b-spline interpolation to correct for motion and magnetic susceptibility interactions. Temporal misalignment between different slices of the functional data (acquired in interleaved Siemens order) was corrected following SPM slice-timing correction (STC) procedure (Sladky et al., 2011), using sinc temporal interpolation to resample each slice BOLD timeseries to a common mid-acquisition time. Potential outlier scans were identified using ART (Whitfield-Gabrieli et al., 2011) as acquisitions with framewise displacement above 0.5 mm or global BOLD signal changes above 3 standard deviations (Nieto-Castanon, 2022; Power et al., 2014), and a reference BOLD image was computed for each subject by averaging all scans excluding outliers. Functional and anatomical data were normalized into standard MNI space, segmented into grey matter, white matter, and CSF tissue classes, and resampled to 2 mm isotropic voxels following a direct normalization procedure (Calhoun et al., 2017; Nieto-Castanon, 2022) using SPM unified segmentation and normalization algorithm (Ashburner, 2007; Ashburner & Friston, 2005) with the default IXI-549 tissue probability map template. Finally, functional data were smoothed using spatial convolution with a Gaussian kernel of 12 mm full width half maximum (FWHM).

*Functional Connectivity Denoising*

Functional data were denoised using a standard denoising pipeline (Nieto-Castanon, 2020b) including the regression of potential confounding effects characterized by white matter timeseries (5 CompCor noise components), CSF timeseries (5 CompCor noise components), motion parameters and their first order derivatives (12 factors) (Friston et al., 1996), outlier scans (below 8 factors for task, below 6 factors for rest) (Power et al., 2014), session effects and their first order derivatives (2 factors for rest, 10 factors for task), and linear trends (2 factors) within each functional run, followed by high-pass frequency filtering of the BOLD timeseries (Hallquist et al., 2013) above 0.001 Hz. CompCor (Behzadi et al., 2007; Chai et al., 2012) noise components within white matter and CSF were estimated by computing the average BOLD signal as well as the largest principal components orthogonal to the BOLD average, motion parameters, and outlier scans within each subject's eroded segmentation masks.

*Diffusion Map Embedding*

Diffusion map embedding employs a random walker to approximate the likelihood of transition between nodes, illuminating the local geometry of the normalized angle matrix. This approach belongs to the family of graph Laplacians (Coifman & Lafon, 2006). It is derived from the equivalence of the distance between points in the low-dimensional embedding space and the diffusion distance between probability distributions at those points. As a result, a transition matrix is generated, across which a Markov chain is run forward in time, linking the local geometries into a set of global axes. These global axes, commonly called gradients in neuroimaging practice, represent the eigenvectors of the diffusion map embedding.

*Representational Similarity Analysis Gradients*

The Representational Similarity Analysis *(*RSA) derived gradients revealed distinct profiles of three gradients that highlighted significant differences in cortical connectivity across different WM operations. Gradient 1 primarily captured the differentiation between sensory-motor and higher-order associative regions, showing the most shifts during maintain vs. replace operations. Gradient 2 delineated the transition between visual and frontoparietal areas, with notable changes during maintain vs. suppress, indicating alterations in sensory integration and executive control. Gradient 3, which captured the differentiation in connectivity between frontoparietal control regions and default mode regions, showed marked differences during maintain vs. clear operations. These distinct gradient profiles underscore the dynamic reorganization of brain connectivity as individuals engage in different WM tasks, with each gradient reflecting specific aspects of neural integration and segregation.

The Posterior Cingulate Cortex (PCC), part of the DM_wm_, exhibited consistent changes across all operation comparisons, particularly in suppress and clear, reflecting its involvement in transitioning between externally and internally focused tasks. The Prefrontal Cortex (PFC) showed changes in replace and suppress conditions. Notable shifts were observed in the V_wm_, especially between maintain and replace, and maintain and suppress. These shifts suggest alterations in how sensory information is integrated during these operations. Notably, the left hemisphere of the V_wm_ showed more changes compared to the right, particularly in areas associated with early visual processing. The SM_wm_ displayed changes predominantly between replace and suppress. Changes in the SM_wm_ were observed more uniformly across both hemispheres. The FPC_wm_ showed marked shifts across all operation comparisons, particularly in replace and suppress, highlighting its role in adapting to different cognitive demands. The right hemisphere regions exhibited more changes in the FPC_wm_, indicating a lateralized function in cognitive control. The DM_wm_ showed gradient shifts between maintain and all other operations, especially suppress and clear. These shifts align with the network's role in self-referential and abstract cognitive processes, with changes more evident in the right hemisphere, pointing to asymmetries in how abstract thinking and internal focus are managed.

*Functional Connectivity Gradients*

Functional Connectivity (FC) Gradient 1 showed shifts in connectivity between primary sensory areas and transmodal regions, especially during maintain vs. replace operations. Gradient 2 highlighted changes in connectivity between visual and cognitive control networks, with differences during maintain vs. suppress, reflecting the integration of sensory information and executive functions. Gradient 3 captured the differentiation between frontoparietal and default mode regions, with larger differentiation during suppress and clear operations.

The PCC exhibited consistent changes across all operation comparisons, particularly in suppress and clear, reflecting its role in switching between external and internal focus. The PFC showed changes in replace and suppress conditions, emphasizing its role in executive functions and cognitive control. The V_wm_ showed shifts between maintain and clear, indicating visual processing and integration changes. The left hemisphere exhibited more changes. The SM_wm_ displayed notable changes between replace and suppress, suggesting alterations in motor coordination and execution. Bilateral changes were observed, similar to the RSA-derived gradients. The FPC_wm_ exhibited shifts across all operation comparisons, particularly in replace and suppress, highlighting its role in adapting to different cognitive demands. Right hemisphere regions showed more changes. The DM_wm_ showed changes between maintain and all other operations, especially suppress and clear. These changes align with its role in abstract cognitive processes and self-referential thinking. Right hemisphere changes were more, indicating asymmetries in managing abstract thinking.

*Representational Similarity Analysis and Functional Connectivity Gradient Comparisons*

Both RSA and FC-derived gradients showed changes in the V_wm_, particularly between maintain and replace, and maintain and suppress. These changes suggest consistent alterations in sensory integration across different methodologies. Both analyses' left hemisphere exhibited more changes, indicating a consistent lateralization pattern. The SM_wm_ displayed notable changes between replace and suppress in both analyses, reflecting its role in motor planning and execution. Bilateral engagement was observed in both RSA and FC-derived gradients. The FPC_wm_ showed marked shifts across all operation comparisons in both analyses, particularly in replace and suppress. This network's involvement underscores its critical role in cognitive control, with right hemisphere regions exhibiting more changes, indicating a consistent lateralized function in cognitive control. Both analyses revealed changes in the DM_wm_ between maintain and all other operations, especially suppress and clear. These shifts align with its involvement in abstract cognition and self-referential thinking, with right hemisphere changes more in both analyses, highlighting asymmetries in abstract cognitive processing.

While the general patterns of change were similar, the magnitude and specificity of changes varied between RSA and FC-derived gradients. For instance, the RSA-derived gradients showed more changes in the PCC during the suppression operation, whereas FC-derived gradients highlighted distinct changes across different operations. Although both analyses revealed right hemisphere dominance in certain networks, the extent of these asymmetries varied. The RSA-derived gradients showed more changes in the right hemisphere DM_wm_, whereas FC-derived gradients indicated a broader distribution of changes across both hemispheres in other networks. The FC-derived gradients highlighted additional changes in regions such as the inferior frontal gyrus and lateral temporal cortex, which were less prominent in the RSA-derived gradients.

*Resting-State Functional Connectivity Gradients*

The resting state gradients revealed functional differentiation running from sensory-to-transmodal (gradient 1), visual-to-insula (gradient 2), and somatomotor-to-insula (gradient 3). Together, these gradients describe functional discrimination between sensory modalities and across levels of the cortical hierarchy (i.e., sensory processing, attentional modulation, and higher-order cognition). Regional gradient values reflect the similarity of connectivity profiles concerning that axis (e.g., two regions with similar gradient 1 values exhibit a similar distribution of rs-FC along the sensory-transmodal axis). Individual embedding solutions for each individual were aligned to the group-level embedding via Procrustes alignments by operation. While both resting-state gradients and FC operation gradients revealed variation in connectivity patterns, they exhibit distinct characteristics compared to the WM operation gradients.

The PCC and PFC showed WM operation-specific changes in the FC operation gradients, with the PCC particularly affected during suppress and clear conditions and the PFC during replace and suppress. These regions may reflect changes in executive functions and cognitive control in FC gradients, while resting-state gradients showed stable connectivity profiles, indicating intrinsic functional hierarchies.

In the V_wm_, FC operation gradients showed changes between maintain and clear conditions, with left hemisphere regions particularly affected, reflecting WM operation-related visual processing and integration changes. Resting-state gradients, however, depicted a stable visual-to-insula axis (gradient 2), indicative of intrinsic connectivity patterns without WM operation-induced difference.

The SM_wm_ exhibited WM operation-induced shifts between replace and suppress conditions in the FC operation gradients, emphasizing motor planning and execution. These changes were bilateral, mirroring the intrinsic somatomotor-to-insula gradient (gradient 3) observed in resting state, which remains stable without WM operation influences.

The FPC_wm_ in the FC operation gradients showed differences across all WM operations, especially replace and suppress. Right hemisphere regions were more affected, consistent with lateralized cognitive functions. Resting-state gradients described this network along the sensory-to-transmodal axis (gradient 1), emphasizing stable higher-order cognitive functions without WM operation-specific modulation.

The DM_wm_ displayed marked changes in the FC operation gradients between maintain and all other conditions, notably suppress and clear. These changes were predominantly in the right hemisphere. Resting-state gradients, on the other hand, showed a consistent transmodal connectivity pattern without such WM operation-driven variations.

*Resting-State vs. Task-Based Metrics*

We conducted a series of regression analyses to evaluate whether resting-state brain organization metrics provide additional predictive power compared to task-based brain organization in explaining thought control problems. The primary goal was to compare and assess whether resting-state metrics improved predictive variance beyond task-based RSA and connectivity metrics. Resting-state metrics corresponding to dispersion and eccentricity were paired by type and _wm_network (e.g., maintain DM_wm_ eccentricity paired with rest DM_wm_ eccentricity) and added to the task-only models to assess the additional predictive power. Nested F-tests were used to determine whether adding resting-state metrics significantly improved model performance. The change in Adjusted R², *F*, and p-values were reported to evaluate the contribution of resting-state metrics.

Analyses revealed that resting-state metrics rarely contributed meaningful improvements in predictive power. Adjusted R² changes were typically small or negligible, and F-tests did not reach significance in most comparisons (see **Supplemental Table 5**).. Task-based metrics generally remained the primary predictors, with resting-state metrics providing little unique explanatory variance or sometimes slightly reducing model fit. Similar patterns emerged for both functional connectivity and RSA metrics, indicating that neural organization during the WM operations more strongly accounts for individual differences in thought control difficulties than resting-state organization.

**Supplemental Figure 1**

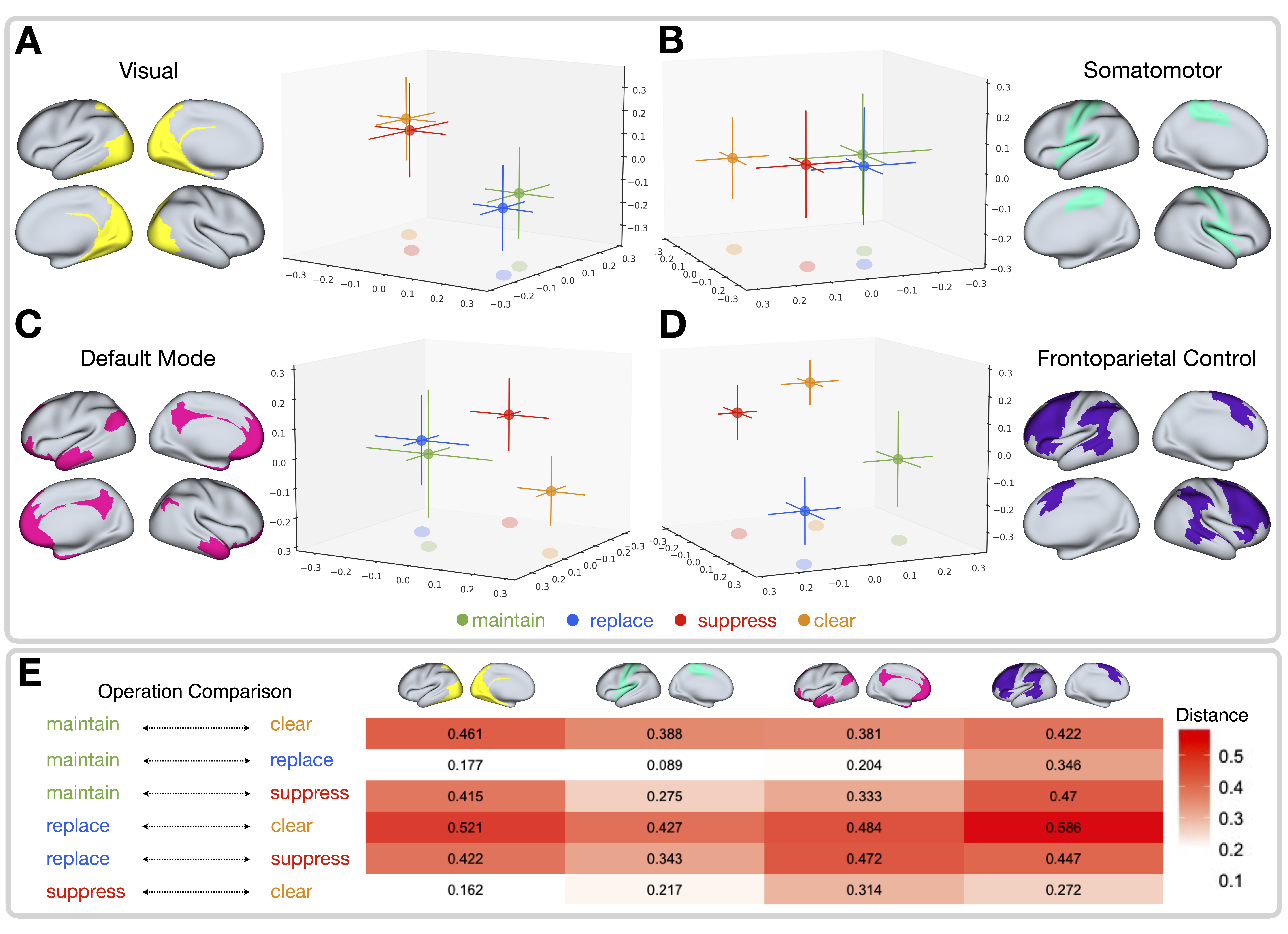
Modified from DeRosa et al. (2024): **A–D**, Within network operation MDS profiles, **A**, Visual _wm_Network, **B**, Somatomotor _wm_network, **D**, Frontoparietal Control _wm_Network, **E**, Default Mode _wm_Network. Single points represent the mean, and bars represent the 95% confidence interval across the MDS representational patterns between trial vectors across the 288 trials (72 for each operation) for each parcel within a given network and colored by operation (maintain, green; replace, blue; suppress, red; clear, orange). **E**, Distances between the pairwise mean MDS pattern across each operation by network. Notes: For a full distribution of the MDS data points that comprise each operation by network, see:

<https://rpubs.com/jakederosa123/Network_Operation_MDS_Full_Distributions>

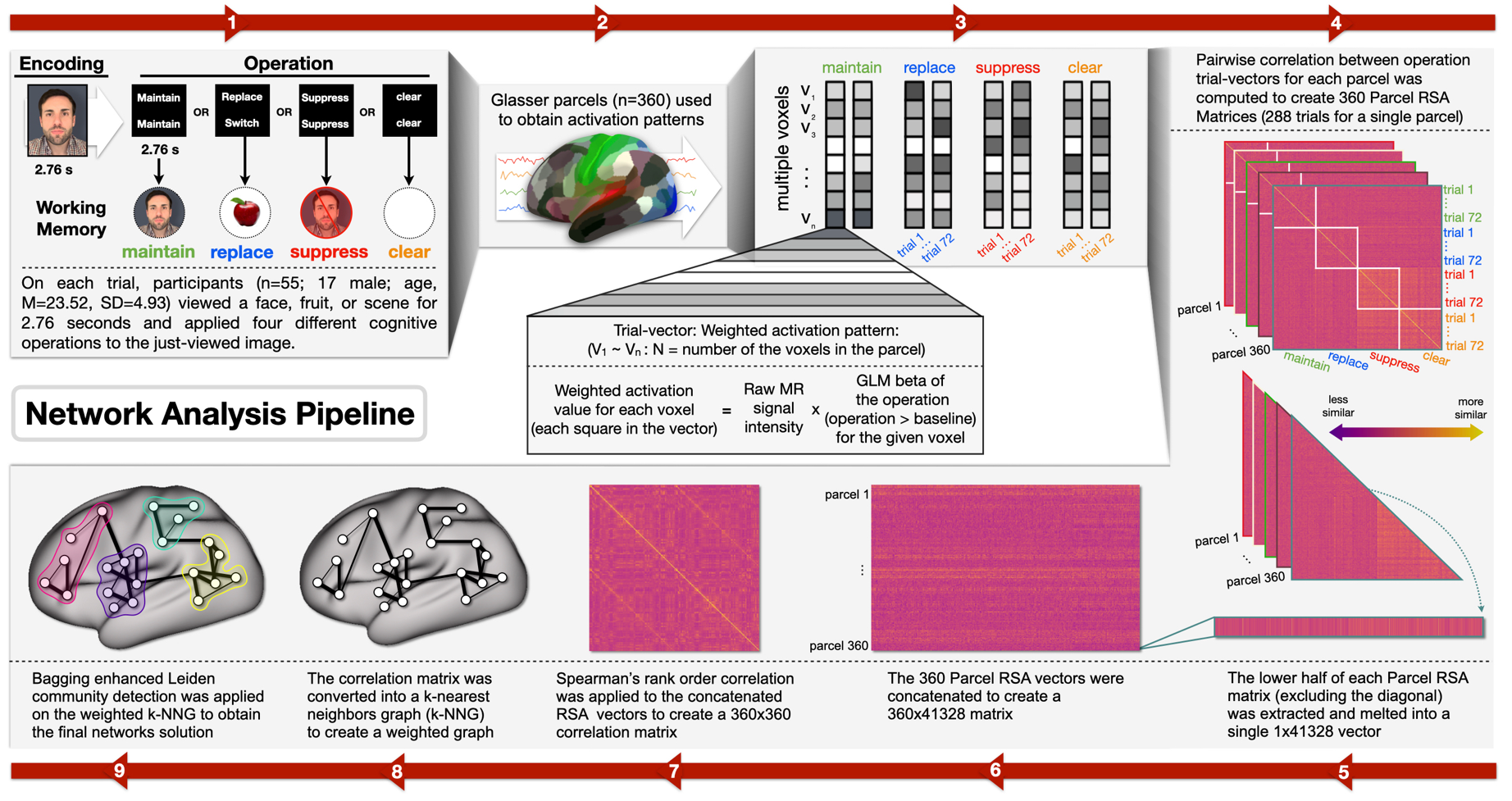
**Supplemental Figure 2**

**1.** DeRosa et al. (2024) outline of the steps to obtain the four representational brain networks. 1) On each trial, participants viewed a picture (a face, fruit, or scene) for 2.76 seconds (s). A screen appeared for 2.76 s indicating which one of four different cognitive operations (*maintain*, *replace*, *suppress*, *clear*) should be applied to the just-viewed image. 2) The Glasser parcellation (Glasser et al., 2016) was used to extract 360 parcels covering a whole brain (180 parcels/hemisphere) for each subject. 3) Activation patterns across all trials (i.e., 72 trial vectors for each operation and 288 trial vectors in total) were computed for each parcel. Trial vectors are weighted activation patterns that consist of multiple voxels (V_1_ ~ V_N_: N = number of voxels in the parcel). The weighted activity value for each voxel equals the raw MR signal intensity multiplied by the GLM beta of the operation (operation > baseline) for the given voxel. The term "raw MR signal" refers to the MR signal intensity extracted from time points at which the operation was performed (including 6TR at the onset of the operation, plus an additional 10TR shift). 4) Parcel RSA similarity matrices were then averaged across all subjects. 4) Pairwise correlation between operation trial vectors for each parcel was computed to create 360 Parcel RSA Matrices (288 trials for a single parcel). 5) The lower half of each Parcel RSA matrix was extracted and melted into a single 1x41328 vector. 6) These 360 Parcel RSA vectors are concatenated to create a 360x41328 matrix. 7) Spearman’s rank order correlation is then applied to the concatenated RSA matrices to create a 360x360 correlation matrix. 8) The 360x360 correlation matrix was converted into a weighted k-nearest neighbor graph (k-NNG) to generate the graph to derive our final representational networks. Weights for the k-NNG were obtained by calculating the Hadamard distances based on the overlap of neighbors (parcels) that represent the relationship between the pairwise similarity of the 360 parcels RSA patterns and their k-nearest neighbors. 9) Bagging enhanced Leiden community detection was applied on the weighted k-NNG to partition the graph to obtain our final (four) network solution.

*Correlation Analysis of WM-Control and Resting-State Neural Metrics*

Supplemental Figures 3-9 (below) present the Spearman rank-order correlations examining the relationship between WM-Control and resting-state neural metrics investigated in this study. These metrics include WM-operation classifier accuracy and measures of WM network organization. The figures illustrate the strength and direction of these associations, with color bars providing a visual representation of the magnitude of the relationships.

**Supplemental Figure 3**

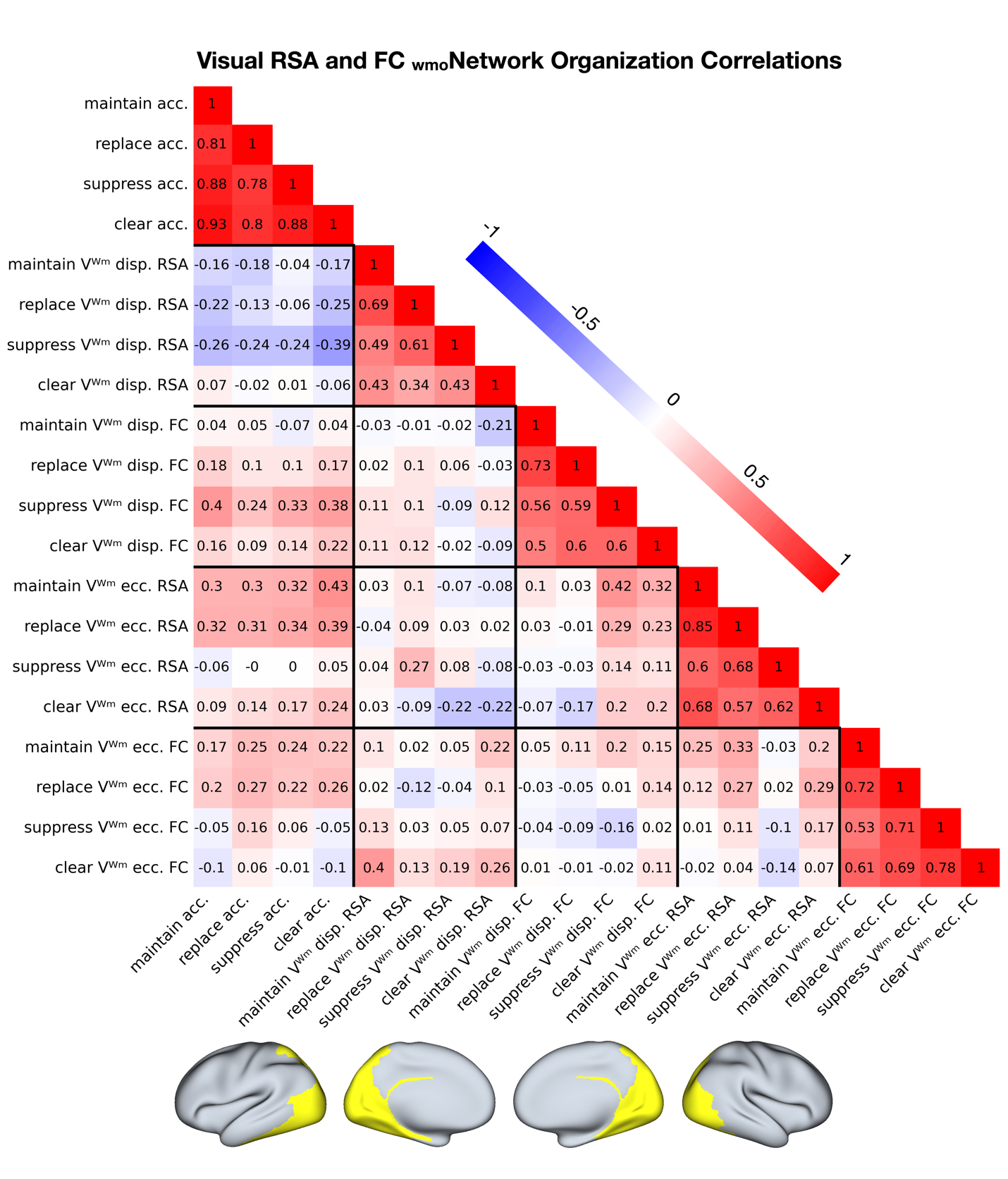

**Supplemental Figure 4**

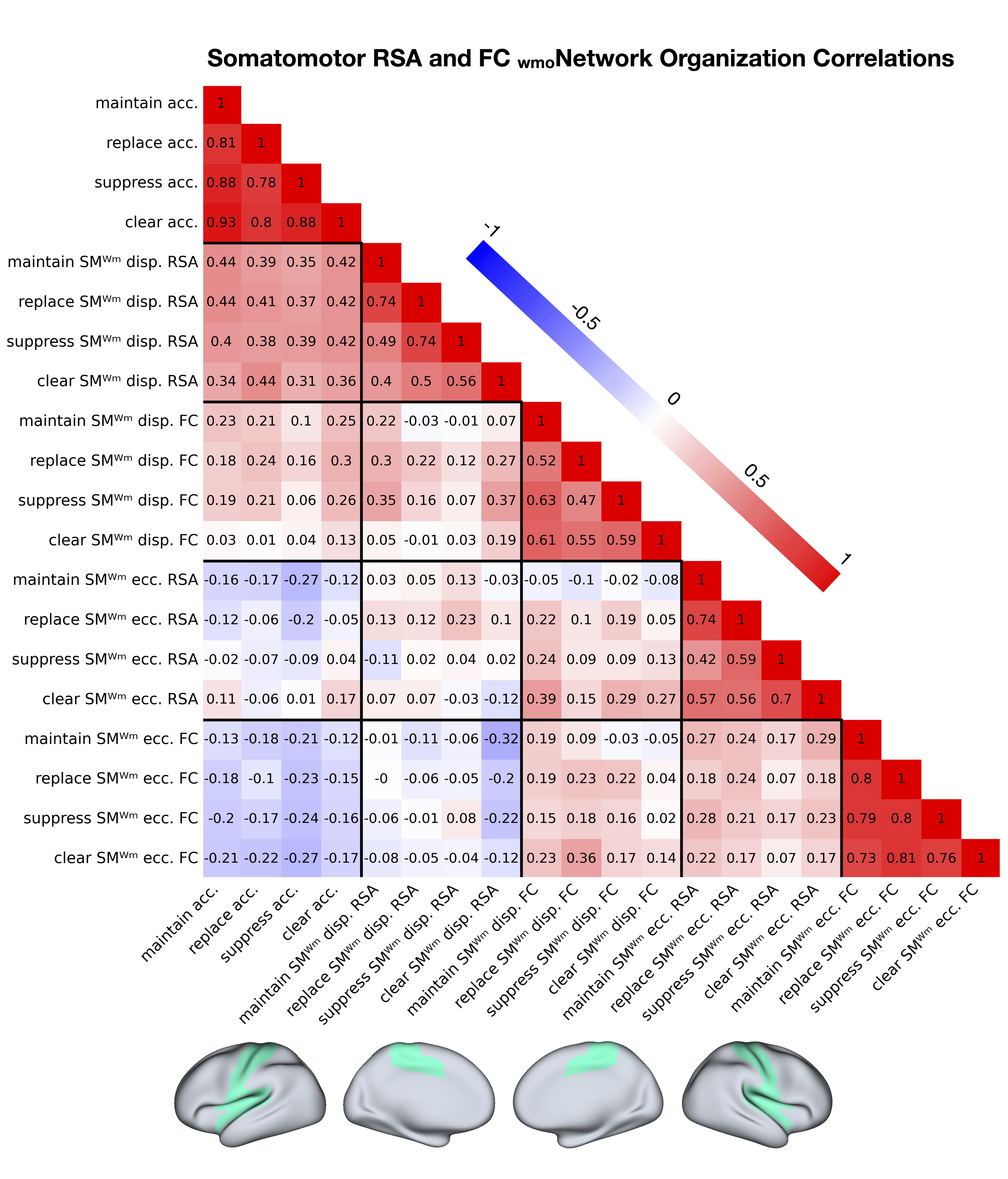

**Supplemental Figure 5**

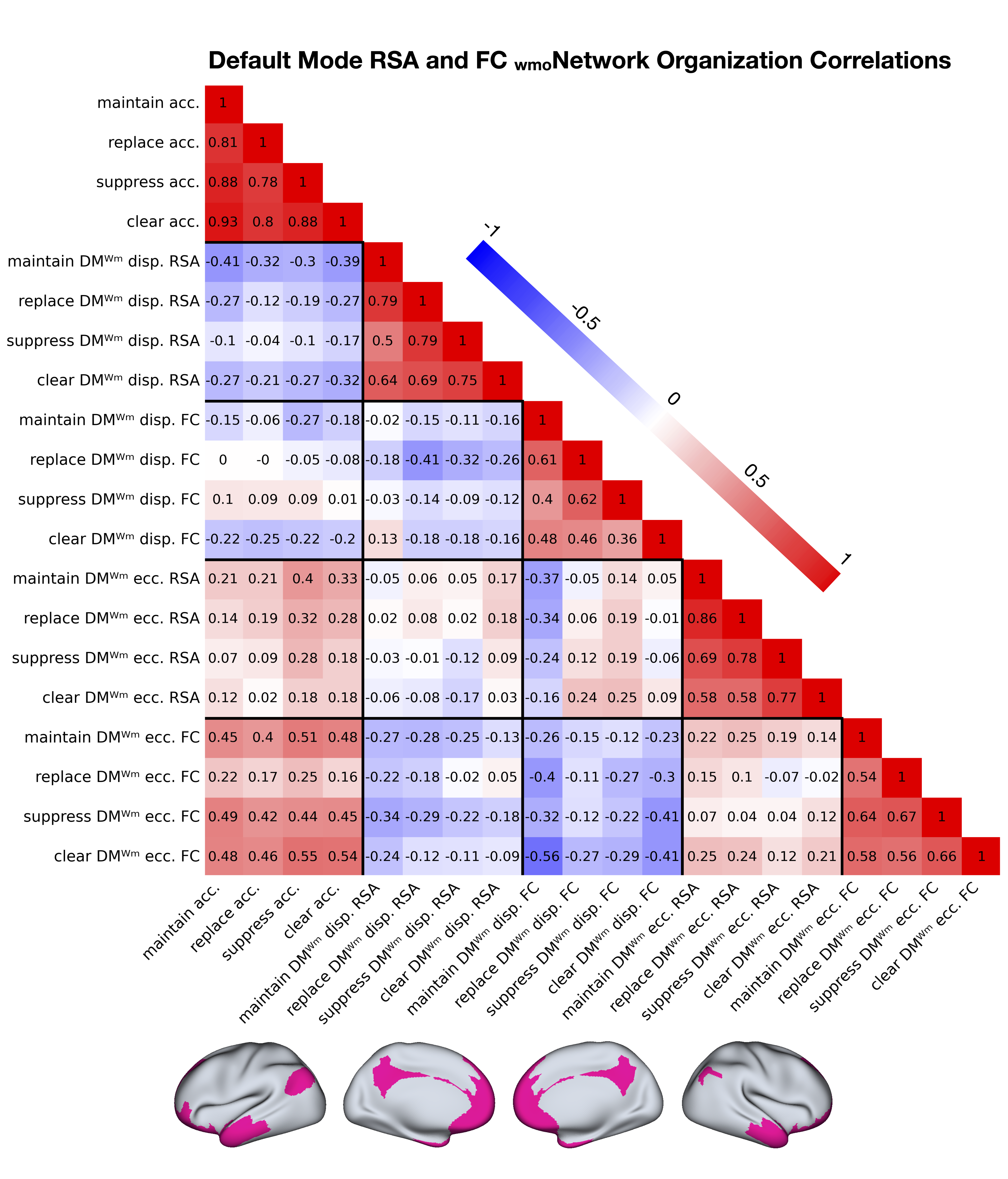

**Supplemental Figure 6**

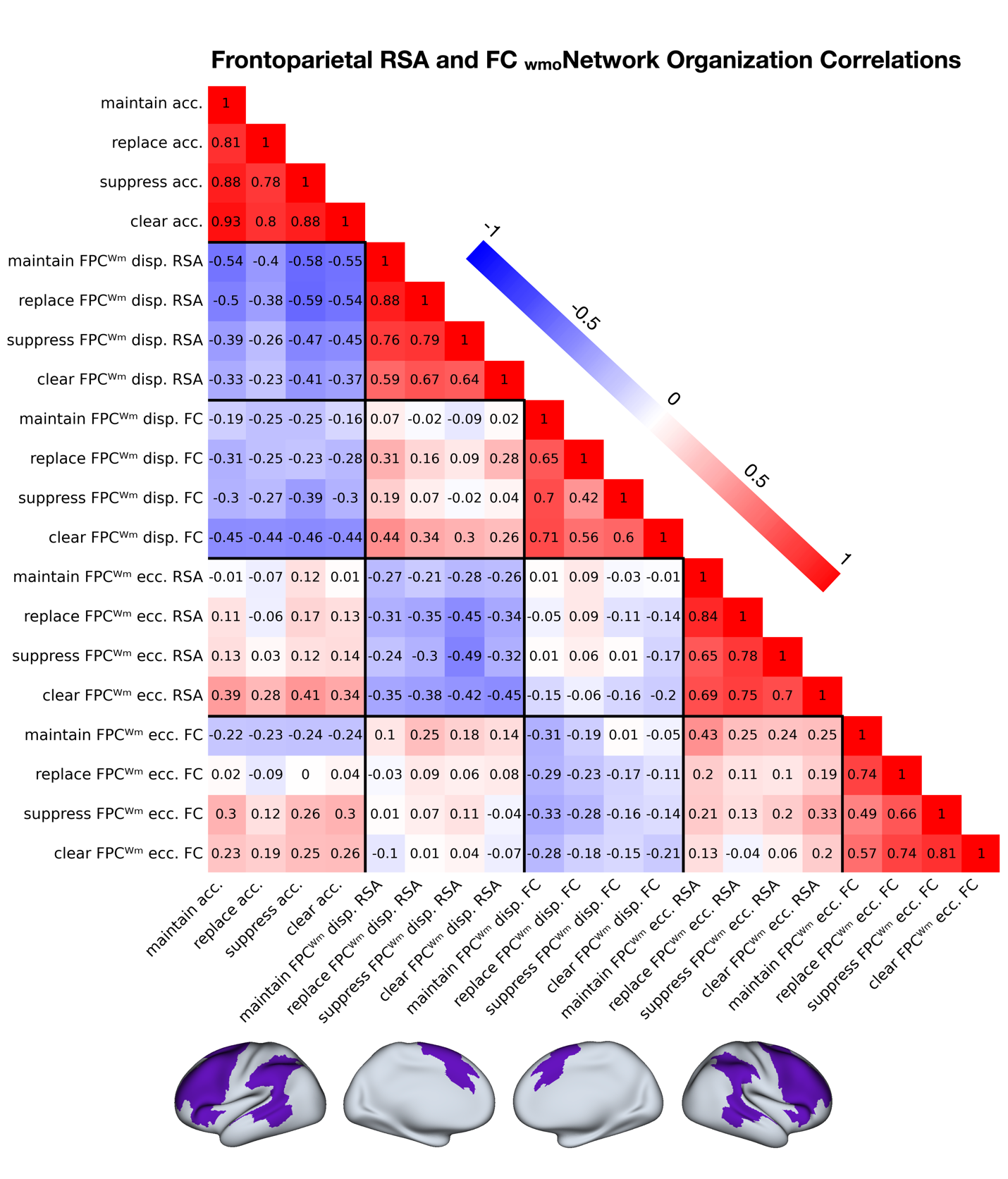

**Supplemental Figure 7**

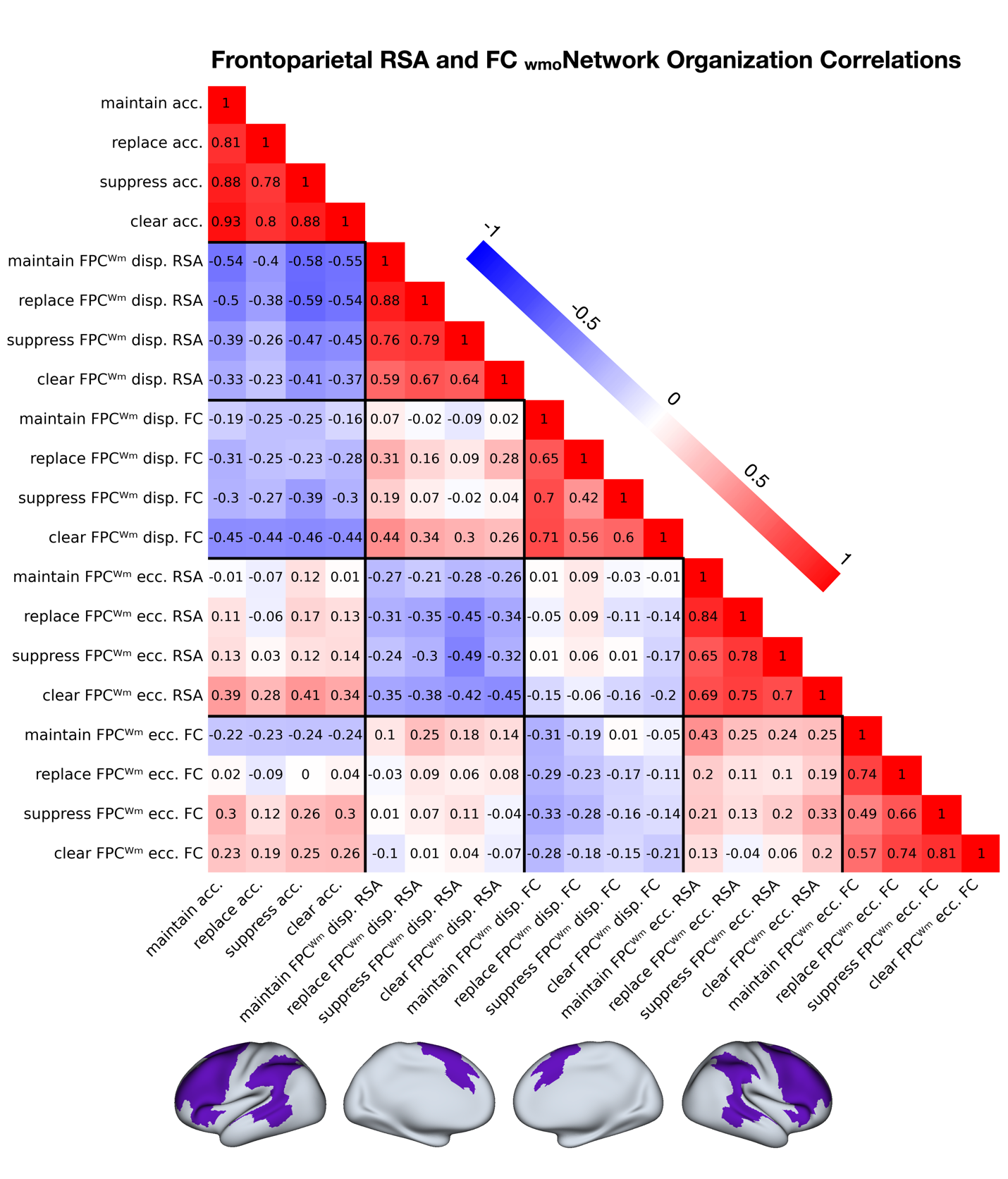

**Supplemental Figure 8**

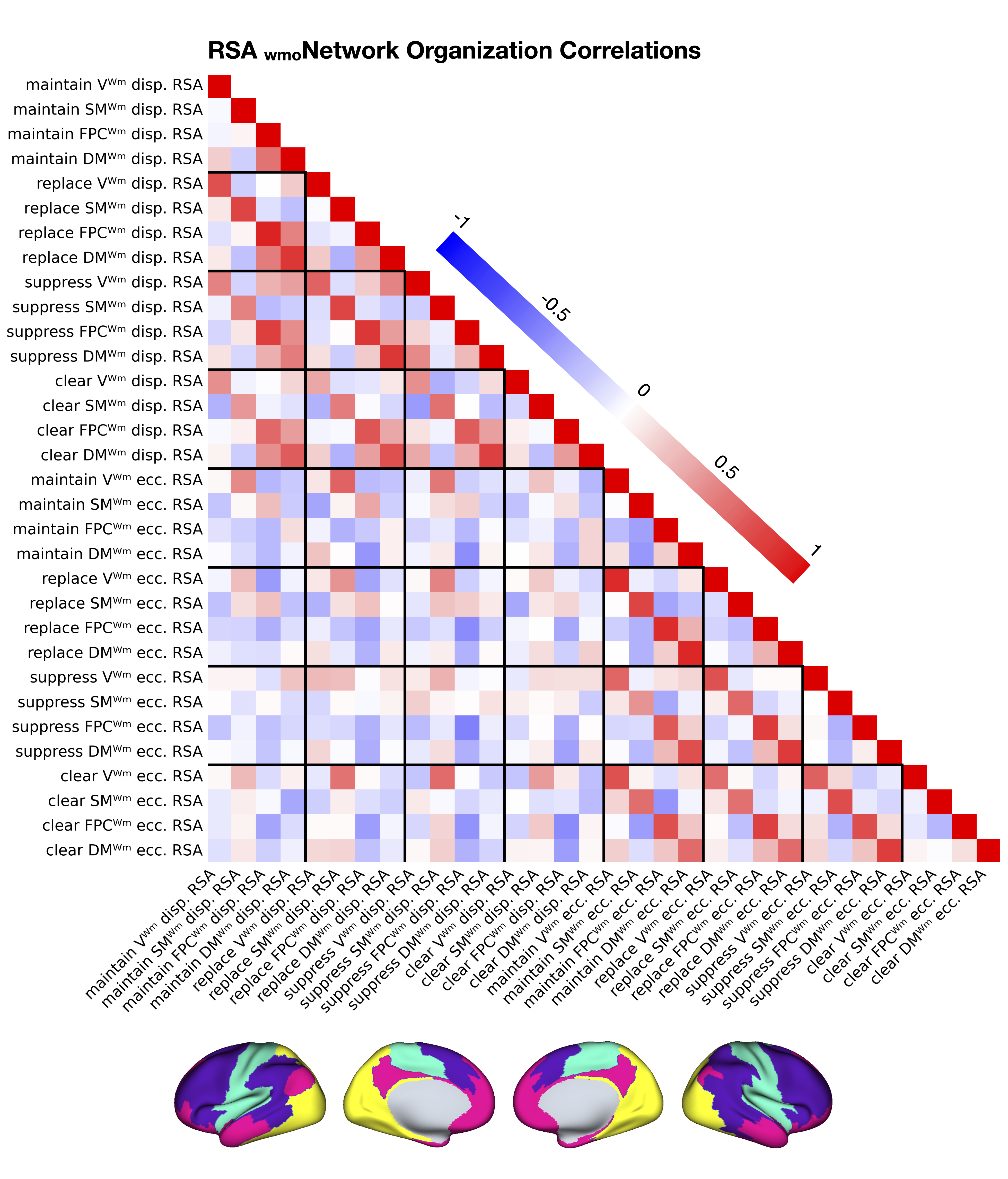

**Supplemental Figure 9**

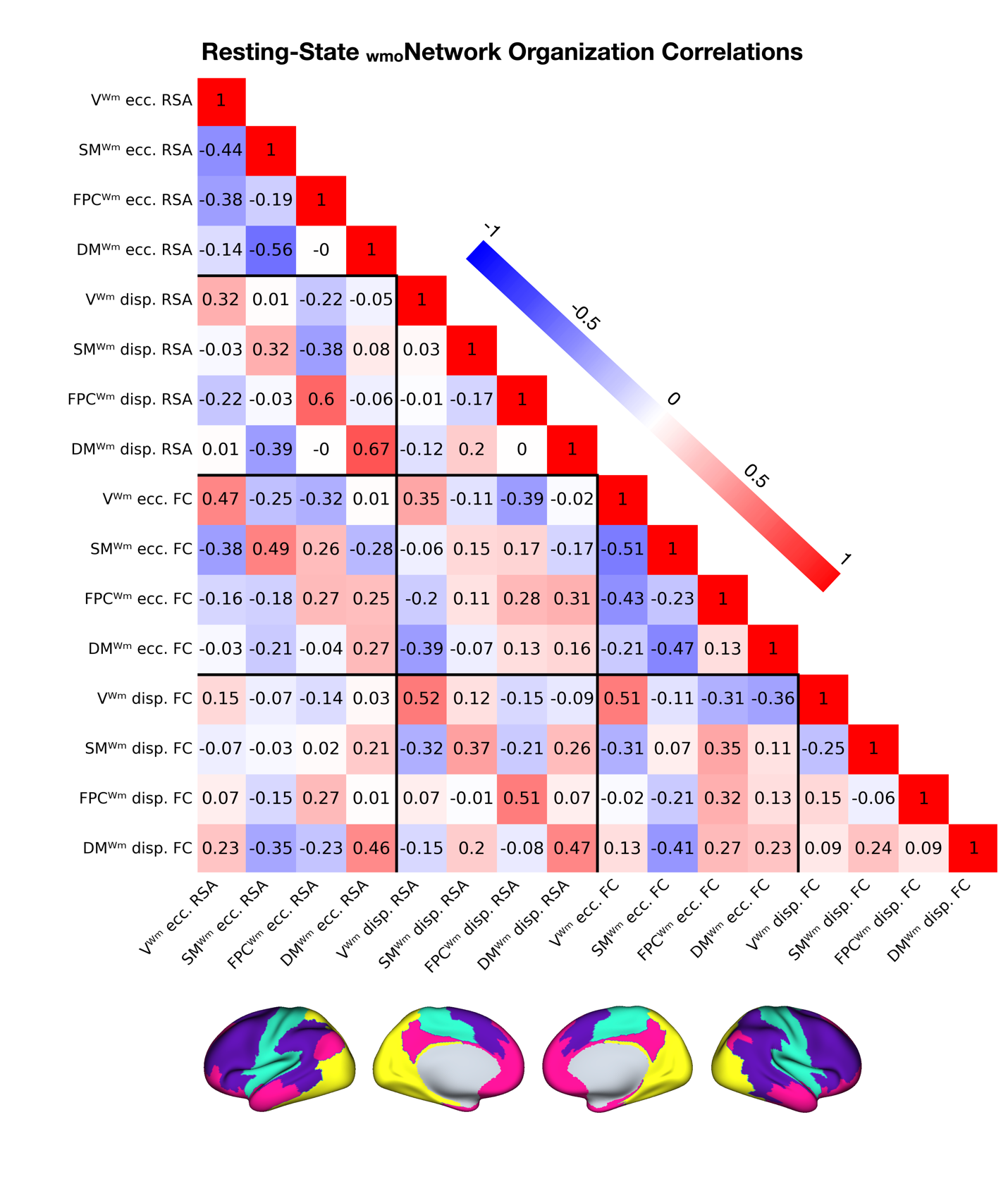
